# BoYueGRN: Zero-shot causal discovery of directed gene regulatory networks from single-cell transcriptomes via amortized inference over synthetic structural causal models

**DOI:** 10.64898/2026.08.15.745056

**Authors:** Jingyue Wu, Ying-Qiang Shen

## Abstract

Gene regulatory network (GRN) inference from single-cell RNA-seq conventionally relies on per-dataset optimization. Existing tools must be refit for every new dataset, and the majority fail to infer causal regulatory directions. Here we present BoYueGRN, an amortized causal discovery framework trained exclusively on 10,000 synthetic structural causal models. For any unseen dataset, a single forward pass returns edge probabilities and regulatory directions, while TF-centric sliding windows with asymmetric fusion extend this fixed-size model to full-transcriptome coverage. BoYueGRN demonstrates strong zero-shot performance across BEELINE benchmarks. On two independent genome-wide CRISPRi Perturb-seq screens, directional accuracy on retained edges reaches 0.86 and 0.95. Reconstructed cell-type- and stage-specific GRN dynamics across five diseases spanning more than 270,000 cells yield experimentally testable biological hypotheses. BoYueGRN reframes directed GRN inference as a train-once, reuse-across-datasets paradigm. By decoupling network reconstruction from per-dataset optimization, this paradigm opens the door to systematic, atlas-scale mapping of regulatory dynamics across human diseases.

## Introduction

Cellular states and functions are orchestrated by GRN, in which transcription factors activate or repress target genes along directed edges[1]. Dissecting these directed relationships is therefore a prerequisite for mechanistic interpretation of cell identity, differentiation, and disease. Single-cell RNA sequencing now provides transcriptomic measurements at single-cell resolution and at a scale of millions of cells, raising the prospect of reconstructing GRNs directly from steady-state single-cell data[2, 3]

A wide range of methods has been proposed to infer GRNs from single-cell transcriptomes, yet systematic evaluations indicate that expression-based reconstruction remains only partially reliable. Early approaches such as ARACNE, CLR, and PIDC score pairwise associations using mutual information or multivariate information measures adapted to single-cell sparsity[4–6]. GENIE3 and its scalable variant GRNBoost2 regress each gene on all remaining genes[7, 8], whereas Gaussian graphical models and shrinkage partial-correlation methods estimate conditional dependencies among genes[9, 10]. A second generation of methods—SCENIC, SCENIC+, CellOracle, FigR, and LINGER—incorporates TF-binding priors from motif databases, ChIP-seq evidence, or paired chromatin accessibility[11–15]. More recent deep-learning models infer regulatory interactions directly from expression profiles[16–19], and pretrained foundation models such as Geneformer, scGPT, and RegFormer offer transferable single-cell representations[20–22]; expression-driven transformer GRN methods such as STGRNS and GeneCompass likewise remain per-dataset or per-corpus learners rather than amortized, train-once directed inference as pursued here. Nevertheless, community assessments of bulk expression profiles have concluded that reliable network inference from gene expression remains an unsolved problem[23, 24], and on the scRNA-seq-oriented BEELINE benchmark most algorithms perform only marginally above random on real networks[25].

Beyond accuracy, three structural limitations constrain the utility of existing tools. The first is coverage: methods whose computational cost grows quadratically with gene number are typically confined to a few hundred highly variable genes, excluding the majority of low- and mid-expression disease-relevant genes. The second is directionality: most tools emit only undirected associations, and prior-knowledge methods can orient edges only where ChIP-seq or motif evidence exists, which is incomplete for most cell types and TFs. The third is the fitting strategy: current methods re-optimize parameters separately for each dataset, which is slow and unstable when genes vastly outnumber cells, and they are designed for a single snapshot rather than for disease progression resolved across stages and cell types. A method that simultaneously delivers genome-wide coverage, regulatory direction, and train-once-reuse-everywhere efficiency remains an unmet need.

Amortized causal discovery offers precisely such a strategy by replacing per-dataset optimization with a model trained once on simulated data and then applied to new data in a single forward pass. Identifiability theory establishes that causal direction can be recovered from observational data alone in both linear non-Gaussian and nonlinear additive noise models[26, 27], and structure learning can be formulated as a continuous optimization problem[28]. These results transfer naturally to scRNA-seq because cell-to-cell variation within a single steady-state experiment carries statistical structure reflecting the underlying regulation, obviating the need for time series or perturbation measurements[3]. ACD on time series and AVICI on synthetic and SERGIO-based semi-synthetic expression data have demonstrated this paradigm[29–31], and the more recent CDFM confirms that zero-shot transfer to real data can succeed when the synthetic distribution spans a sufficiently broad mechanistic space[32]. Sim-to-real transfer is well established in other fields, for example domain randomization in robotics[33]. However, whether it can be applied directly to GRN inference remains unclear. That is because GRN inference confronts distinct challenges: a high-dimensional space of tens of thousands of genes, extremely sparse expression with dropout noise, directed TF-to-target regulation, and scale-free hub topology. To date, no amortized causal discovery framework has been tailored to these constraints, leaving a critical methodological gap for scalable, directed GRN inference from observational single-cell data.

Here we present BoYueGRN, an amortized causal discovery framework for directed GRN inference from scRNA-seq that closes this gap. The model is trained entirely on 10,000 synthetic expression datasets generated by structural causal models (SCMs) whose topology, noise mechanisms, and expression statistics are aligned with real GRNs; for any new dataset, a single forward pass returns edge probabilities and regulatory directions, and TF-centric sliding windows extend this fixed-size model to full-transcriptome coverage. The name BoYue, drawn from the Chinese adage “**b**o guan er **y**ue qu” (coined by the renowned Song dynasty poet Su Shi, meaning “survey broadly, then extract selectively”), captures this design: broad pretraining on synthetic causal models followed by selective inference in a single forward pass. Having seen no real data during training, BoYueGRN already outperforms several mainstream methods at the TF-centric single-window scale and achieves the best AUROC on all six BEELINE datasets under full-genome coverage; on two genome-wide CRISPRi perturbation screens it attains strong directional accuracy. Building on these benchmarks, we apply BoYueGRN to five disease settings spanning more than 270,000 cells, where the model reconstructs cell-type-and stage-specific GRN dynamics, reproduces known pathological regulatory features, and generates experimentally testable hypotheses. BoYueGRN thus reframes directed GRN inference as train-once-reuse-across-datasets, converting network reconstruction from a per-dataset analysis into a reusable capability for disease research.

## Results

### The BoYueGRN framework

BoYueGRN is an amortized framework for directed GRN inference from scRNA-seq, trained exclusively on synthetic SCMs and, once trained, requiring only a single forward pass to produce a directed, genome-wide GRN for any new real dataset (Fig. 1).

**Fig. 1.**
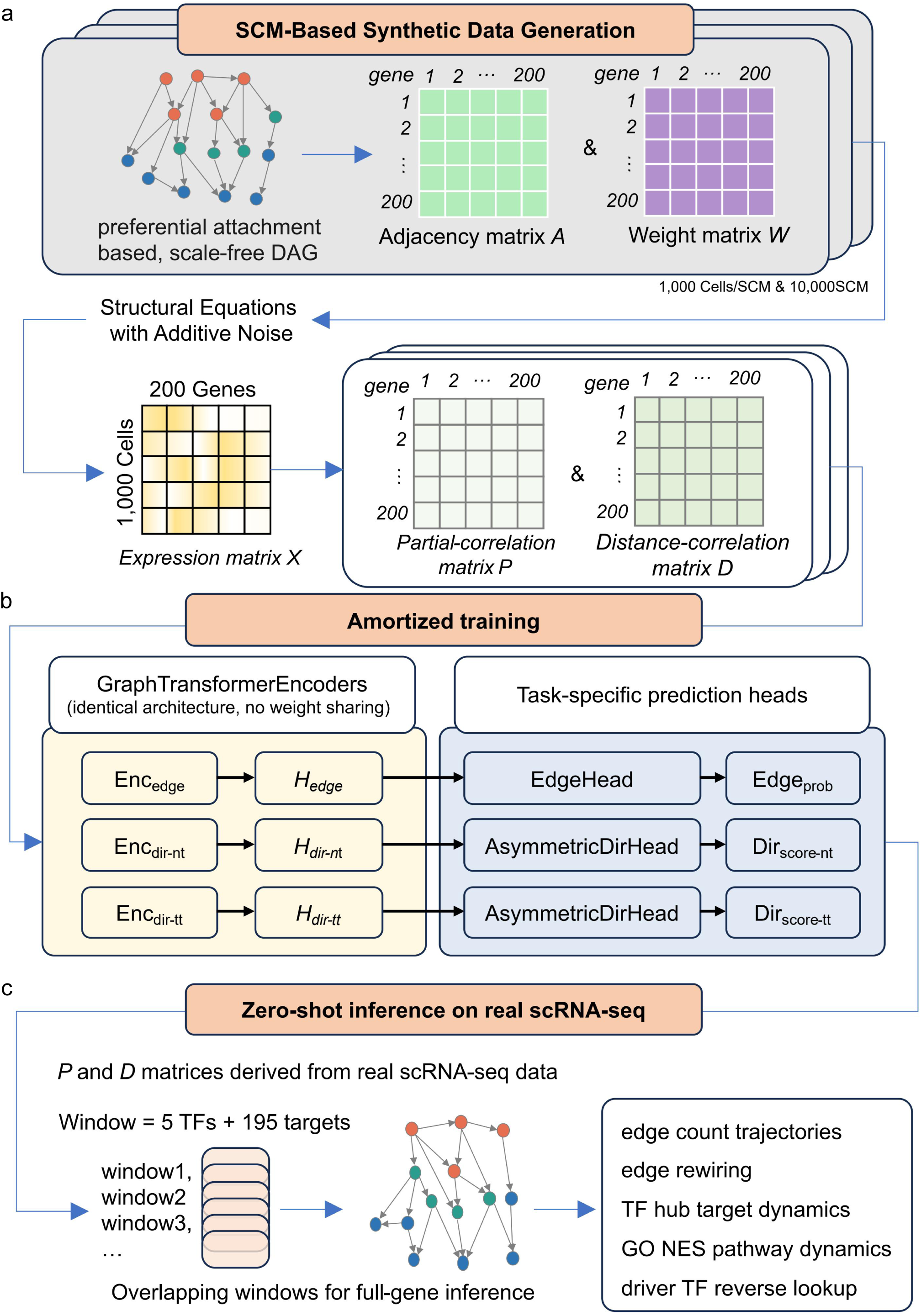
The BoYueGRN framework. a, SCM-based synthetic data generation. Graph topologies are scale-free DAGs grown by preferential attachment (a five-node seed causal chain; each new node samples parents from earlier-indexed nodes with probability proportional to in-degree plus one, guaranteeing acyclicity), yielding the directed adjacency matrix A and the weight matrix W (1,000 cells per SCM; 10,000 SCMs in total). b, Expression simulation and amortized training. Each node’s expression is generated in topological order through structural equations with additive noise (about 50% linear and 50% GELU mechanisms), producing the 200-gene × 1,000-cell expression matrix X, from which the Ledoit–Wolf shrunk partial-correlation matrix P and the distance-correlation matrix D are computed; three GraphTransformer encoders of identical architecture with no weight sharing (Enc_edge, Enc_dir-nt, Enc_dir-tt) feed task-specific prediction heads (EdgeHead producing Edge_prob; AsymmetricDirHeads producing Dir_score-nt and Dir_score-tt). c, Zero-shot inference on real scRNA-seq. P and D matrices derived from real data are scored within TF-centric overlapping windows (5 TFs plus 195 correlation-preselected targets each) that slide across the transcriptome for full-gene coverage; the resulting directed GRN supports five downstream analyses: edge-count trajectories, edge rewiring, TF-hub target dynamics, GO-NES pathway dynamics, and driver-TF reverse lookup.

We first construct the training corpus by simulating 10,000 synthetic expression datasets whose graph topology is generated by preferential attachment and whose expression follows additive noise structural equations (Fig. 1a, b). A five-node seed causal chain is created first, after which each node selects parents from earlier-indexed nodes with probability proportional to in-degree plus one; constraining parents to earlier-indexed nodes guarantees acyclicity by construction and reproduces the scale-free hub structure of real GRNs[34]. Each node’s expression is generated in topological order through additive noise structural equations (roughly 50% linear and 50% GELU mechanisms),

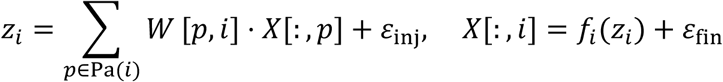

where the magnitude gap between the injection noise ε_inj_ ∼ *N*(0, 0.3^2^) and the final noise ε_fin_ ∼ *N*(0, 2.0^2^) supplies the structural basis for directional identifiability[27], and a log1p transform of the non-negative expression yields the non-negative, sparse, approximately log-normal statistics characteristic of scRNA-seq. Two complementary gene-pair features serve as model inputs— the Ledoit–Wolf shrunk precision matrix ***P* [35]**, capturing linear conditional dependence, and the distance correlation matrix ***D* [36]**, capturing nonlinear dependence—while the directed adjacency matrix **A** is retained as supervision for both the edge-existence and direction tasks (full definitions in Methods).

Three graph Transformer encoders of identical eight-layer architecture then learn node representations from P+D, serving the edge-existence task and two target-typed direction tasks with separately optimized heads (Fig. 1b, Fig. S1). One encoder is dedicated to edge existence; the other two handle the TF-to-non-TF and TF-to-TF direction tasks, respectively. The edge encoder is trained from scratch, while the two direction encoders are warm-started from its weights and fine-tuned independently so that no parameters are ultimately shared. All three use edge-biased attention, superimposing a linear projection of gene-pair statistics onto the standard QKᵀ dot product to couple message passing with pairwise dependence strength. Contrasting weight cosine similarity, parameter distributions, and output embedding-space structure confirms that the task heads learn clearly differentiated representations (Fig. S3). The edge-existence head scores whether an edge exists via a symmetric dot product, whereas the direction head scores regulatory direction via asymmetric source/target projections (Fig. S2),

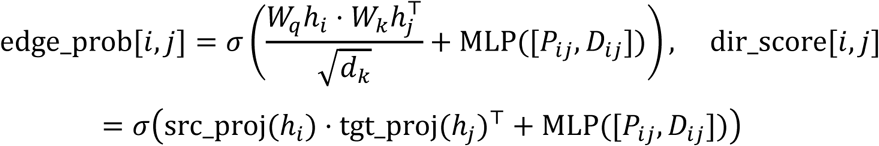

Because the two projections differ, the ordered-pair scores dir_score[*i*, *j*] and dir_score[*j*, *i*] are decoupled by construction, structurally demanding features that distinguish the regulator role from the target role; these features conflict with those needed by the edge-existence head, motivating separate training of the two tasks. The direction head is further split into dual experts by target-gene type: the TT expert is trained only on TF-to-TF ground-truth edges and the NT expert only on TF-to-non-TF edges, and because genome-wide validation on real data shows the TT expert to be superior on both edge types, the deployed pipeline routes every edge to the TT expert (Methods); the matched-expert advantage on synthetic data (Fig. 2e) reflects the synthetic training distribution and does not transfer to real inputs, which is why the deployed routing rule is uniform rather than type-matched. All models are trained only on synthetic data (loss functions in Methods).

**Fig. 2.**
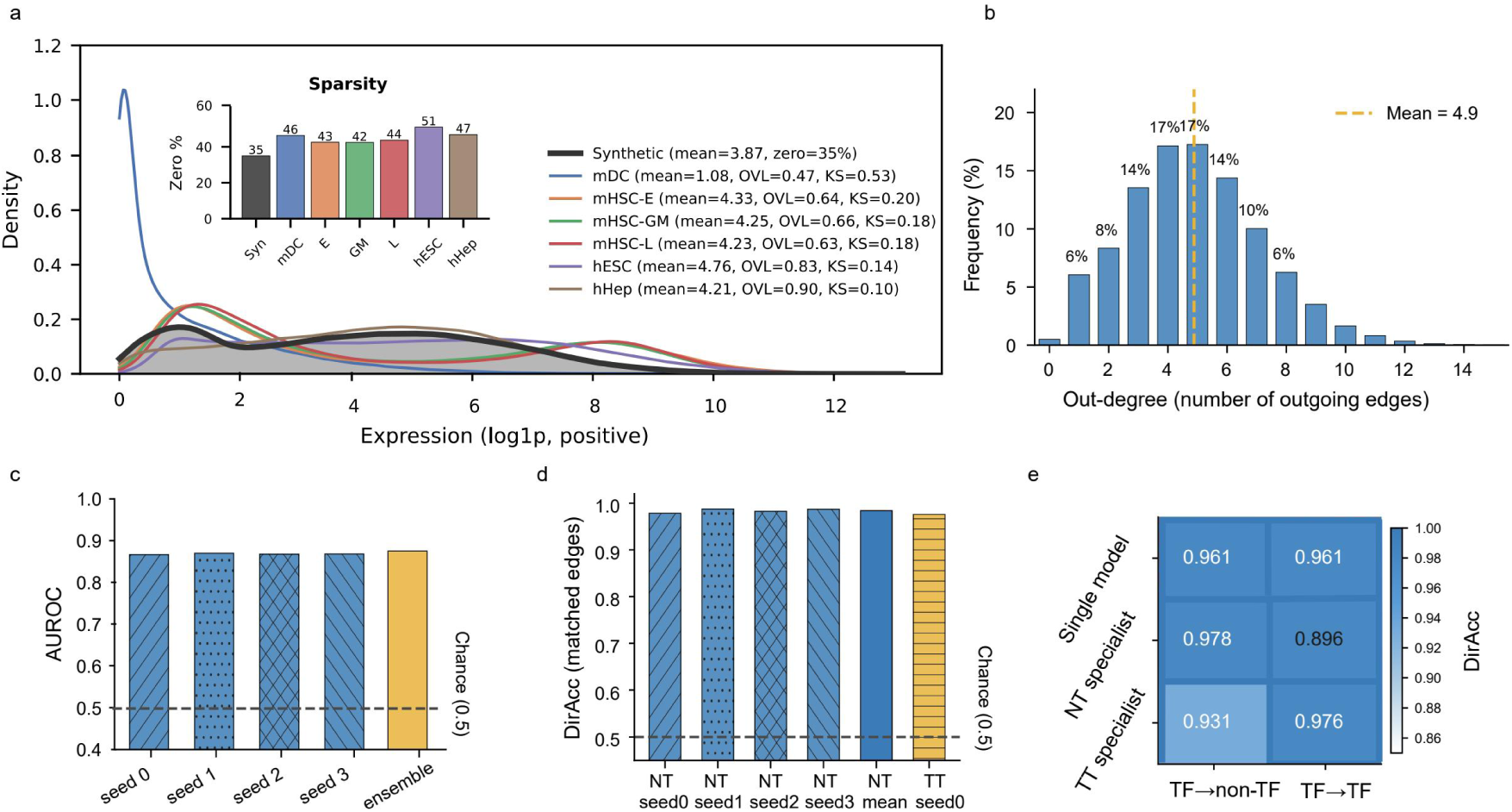
Fidelity of the synthetic training distribution and task learnability. a, Density of positive log1p expression for synthetic data (mean 3.87; 35% zeros) versus the six BEELINE datasets (positive-expression means 1.08–4.76; overlap coefficients 0.47–0.90; KS distances as indicated); inset: sparsity (percentage of zeros) of synthetic and real datasets (42–51%). b, Out-degree distribution of the synthetic networks (mean 4.9, long tail), reproducing the scale-free hub topology of the preferential-attachment generator. c, Edge-prediction AUROC across four independent training seeds (approximately 0.87 each) and after four-seed probability ensembling (0.875). d, Directional accuracy (DirAcc) on matched edge types for the NT expert (four seeds and their mean) and the TT expert on synthetic test data (0.976–0.987). e, Matched versus mismatched expert assignment (DirAcc heatmap): the unspecialized single model achieves 0.961 on both edge types; the NT specialist achieves 0.978 on TF→non-TF edges but 0.896 on TF→TF edges, and the TT specialist achieves 0.976 on TF→TF edges but 0.931 on TF→non-TF edges.

Finally, to achieve full-transcriptome coverage from a model trained and inferred at a fixed G=200-gene scale, TF-centric windows slide across the genome and their predictions are merged by complementary fusion rules (Fig. 1c). Each window contains 5 TFs and 195 target genes (preselected by Pearson correlation and rotated across windows), so each TF is examined alongside its candidate targets in multiple windows. Edge probability is fused by taking the maximum across windows as a recall-oriented strategy, whereas direction score is fused by taking the mean across windows as a multi-window consensus.

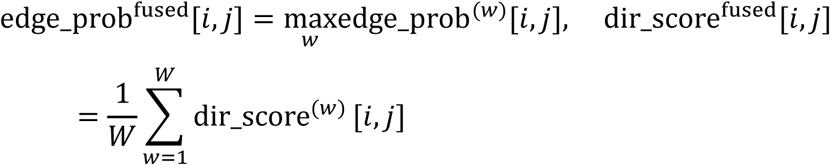

Directed edges are emitted at threshold τ=0.2, and the resulting directed GRN directly supports the five downstream analyses shown on the right of Fig. 1c: edge-count trajectory, edge rewiring, TF-hub target dynamics, GO-NES pathway dynamics, and driver-TF reverse lookup.

We next verify that the synthetic training distribution faithfully mimics real transcriptomes and that the model has learned both target tasks, since alignment between training and inference distributions is a prerequisite for zero-shot transfer.

After log1p transformation, synthetic expression reproduces the qualitative shape of real scRNA-seq—non-negative, sparse (35% zeros, comparable to the 42–51% of the real datasets), and approximately log-normal—and its positive-expression distribution (mean 3.87) closely matches that of five of the six BEELINE datasets (positive-expression means 4.21–4.76; overlap coefficient 0.63–0.90, KS distance 0.10–0.20; Fig. 2a), while the average out-degree of synthetic networks is 4.9 with a long tail extending to 15, reproducing the scale-free hub structure of the preferential attachment generator (Fig. 2b). The edge model yields highly reproducible AUROC values (∼0.870) across four random seeds (Fig. 2c). Averaging predicted probabilities from these four independently trained models during inference mitigates seed-dependent noise and further improves the AUROC to 0.875, indicating that edge-prediction ability is stable and initialization-independent (Fig. 2c). For direction prediction, experts matched to edge type reach DirAcc of 0.976–0.987 on their respective edge types (Fig. 2d). Against an unspecialized single-model baseline, matched experts are superior on their own edge types: 0.978for the NT expert on TF-to-non-TF edges and 0.976 for the TT expert on TF-to-TF edges. By contrast, mismatched combinations systematically degrade (NT expert 0.896 on TF-to-TF edges; TT expert 0.931 on TF-to-non-TF edges) (Fig. 2e). Collectively, validations covering data fidelity, edge learnability, and direction separability support the model’s transferability to real scRNA-seq data.

### Edge and direction prediction benchmarks

We next test whether the edge-prediction and direction-inference abilities established on synthetic data transfer to real scRNA-seq. Zero-shot evaluation is performed on the six BEELINE benchmark datasets (mDC, mHSC-E, mHSC-GM, mHSC-L, hESC, hHep), with ground-truth networks taken from the ChIP-seq-derived networks released with BEELINE, and BoYueGRN is compared against seven established methods (GENIE3, GRNBoost2, Correlation, MutualInfo, PPCOR, GLasso, and dCor) plus a random baseline on identical inputs. Direction prediction is assessed on the same benchmark and further validated against causal ground truth on two genome-wide Perturb-seq screens.

#### Edge-existence prediction

At the TF-centric single-window setting (G=200), BoYueGRN attains the highest AUROC on five of six datasets (mHSC-E 0.749, mHSC-GM 0.760, mHSC-L 0.751, hESC 0.736, hHep 0.604; mean 0.679; Fig. 3a). Beyond overall ranking, the early precision ratio (EPR), the enrichment multiple of ground-truth edges among the top-k predictions, exceeds 2 on four of six datasets, peaking at 2.97 for mHSC-L (Fig. 3b). Extending the model to full-genome coverage via window sliding raises the mean AUROC to 0.760 and keeps EPR above 4 on all six datasets (Fig. 3c–d).

**Fig. 3.**
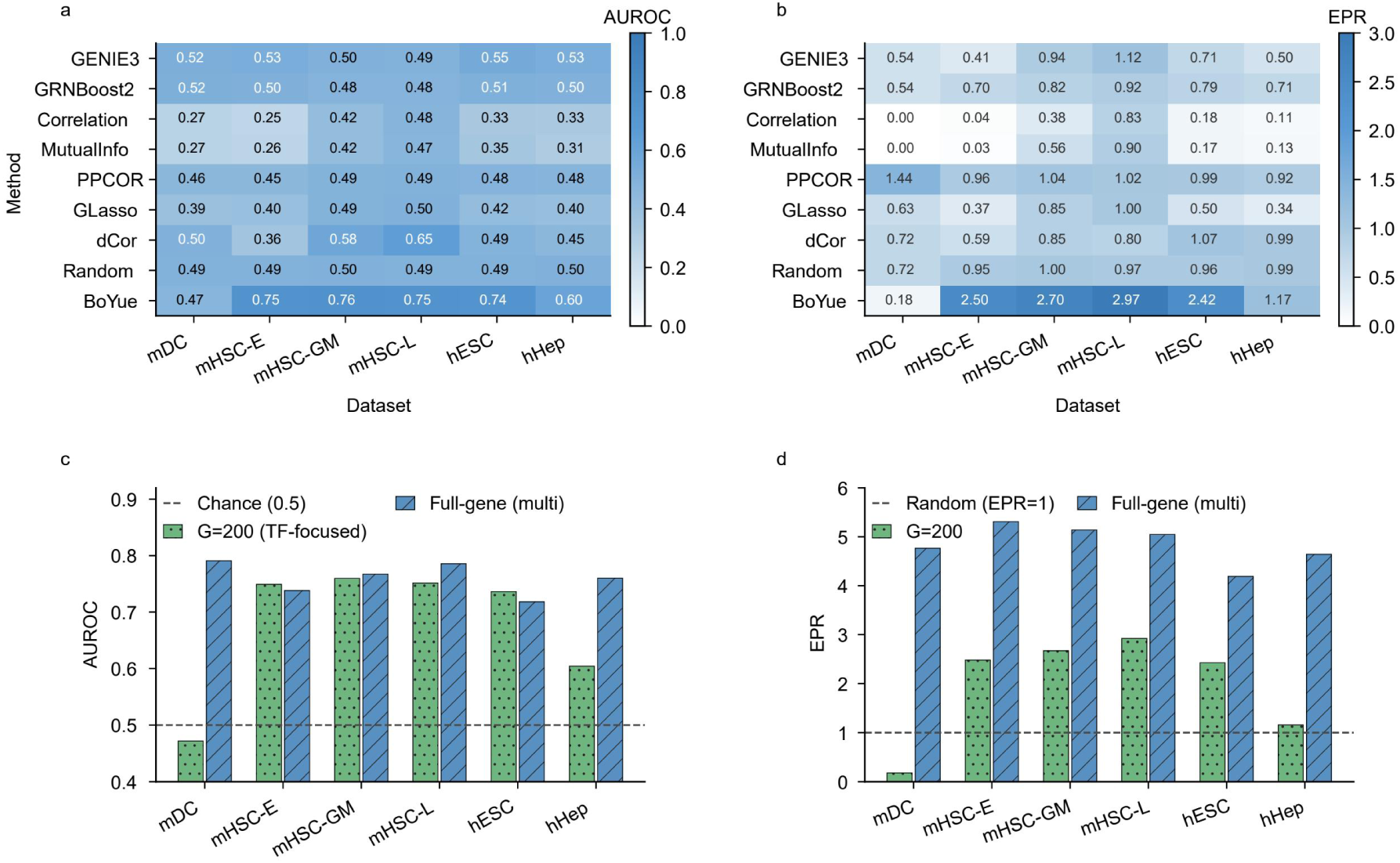
Zero-shot edge prediction on the BEELINE benchmark. a, Single-window (G=200) AUROC of BoYueGRN and seven baseline methods (GENIE3, GRNBoost2, Correlation, MutualInfo, PPCOR, GLasso, dCor) plus a random baseline across the six BEELINE datasets; BoYueGRN attains the highest AUROC on five of six datasets (mHSC-E 0.749, mHSC-GM 0.760, mHSC-L 0.751, hESC 0.736, hHep 0.604), with mDC at 0.47. b, Early precision ratio (EPR) under the same single-window setting (BoYueGRN: 2.50, 2.70, 2.97, and 2.42 on mHSC-E, mHSC-GM, mHSC-L, and hESC). c, AUROC at G=200 versus genome-wide (full-gene, multi-window) coverage; genome-wide inference outperforms all baselines on all six datasets (mean AUROC 0.760), including recovery of the distributionally outlying mDC dataset (0.472 to 0.791). d, EPR at G=200 versus genome-wide coverage, exceeding 4 on all six datasets in genome-wide mode. The hESC values shown incorporate the variance-based rank-deficiency guard (Table 1).

**Table 1.** hESC genome-wide evaluation before and after the rank-deficiency guard.

| Metric | Before | After | Change |
| --- | --- | --- | --- |
| Input genes | 17,735 | 10,000 | — |
| AUROC | 0.501 | 0.718 | +0.217 |
| EPR | 1.99× | 4.19× | $\times 2.1$ |
| DirAcc | 0.698 | 0.841 | +0.143 |

#### Small-sample recovery via multi-window ensembling

The single dataset on which single-window performance initially lags, mDC, illustrates that this is an explainable statistical-estimation problem rather than a model deficiency, and its recovery provides direct evidence for the robustness of multi-window asymmetric fusion. Under the single-window G=200 setting, BoYueGRN’s initial AUROC on mDC is only 0.472 with an EPR of 0.18—the only one of the six datasets that fails to clearly lead the baselines. mDC has only 383 cells, 30 TFs, and 657 ChIP-seq ground-truth edges—the lowest edge count and edge density (≈22 edges per TF) among the four ChIP-seq–annotated datasets, making it both the smallest and, within its ground-truth class, the most sparsely annotated of the six. It is also the only dataset whose expression distribution departs markedly from the rest (positive-expression mean 1.08 vs 4.21–4.76; overlap coefficient 0.47 vs 0.63–0.90; Fig. 2a), so the model operates partly out-of-distribution on it. At the high-dimensional ratio n≈383, p=200, the Ledoit– Wolf shrunk precision matrix degrades and the distance correlation matrix likewise suffers small-sample noise, and the statistical uncertainty of both input features—compounded by this distributional mismatch—jointly suppresses single-window performance. Consistent with a dimensionality effect, ablation shows that mDC’s optimal single-window size is G=150 rather than G=200. When the model is extended to genome-wide coverage, however, the max-fusion integration of 3,990 overlapping windows averages out single-window statistical noise: mDC’s AUROC recovers from 0.472 to 0.791 and its EPR rises from 0.18 to 4.76, a level fully consistent with the other five datasets (EPR 4.19– 5.31). That this recovery holds even for the distributionally outlying dataset underscores that the robustness of multi-window asymmetric fusion extends beyond statistical noise to moderate distribution shift. In genome-wide mode BoYueGRN outperforms both random guessing and all baseline methods on all six datasets.

#### Direction inference and confidence calibration

Using the stronger of the two direction experts (the TT expert), BoYueGRN exceeds the unspecialized scheme on all six BEELINE datasets, with four reaching DirAcc ≥ 0.83 (Fig. 4a). Because ChIP-seq ground truth reflects physical binding rather than causal regulation, we further test direction inference on two independent human Perturb-seq datasets that provide causal ground truth from TF knockdown: HCT116, from the X-Atlas/Orion genome-wide Perturb-seq atlas of colon cancer generated on the FiCS fixation-freezing platform[37], and K562, from the genome-wide CRISPRi screen of Replogle et al.[38].Accuracy falls below the random level (0.5) at the default G=200 input scale, but genome-wide coverage raises it to 0.821 and 0.810, respectively (Fig. 4b). The below-chance single-window values indicate a systematic inversion rather than an absence of signal: at G=200 the window is largely filled with genes unrelated to the perturbed TF, so pairwise direction comparisons are dominated by noise, whereas genome-wide windows pair each TF with its correlation-preselected candidate targets and mean-aggregate direction scores across many windows, which removes the inversion. Per-condition decomposition shows that on retained edges (edge_prob > 0.2) accuracy reaches 0.859 (HCT116) and 0.947 (K562) overall and 0.949/0.958 on the TF-to-TF subset, while shuffled controls return exactly to the random level (Fig. 4c–d). Collectively, direction inference yields both high accuracy and trustworthy prediction confidence. Its probability outputs can therefore be directly deployed for downstream edge filtering and driver-gene ranking.

**Fig. 4.**
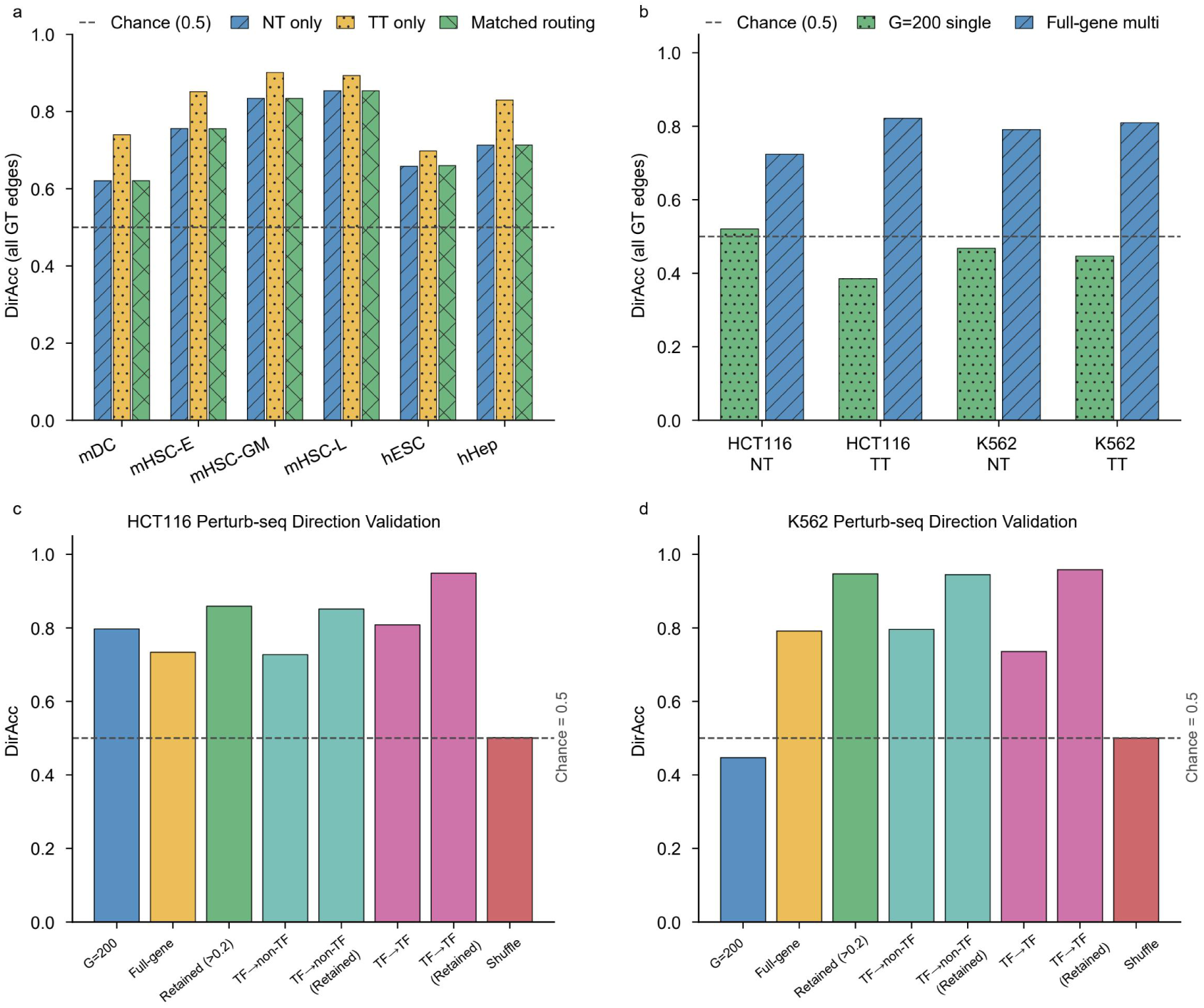
Direction inference on BEELINE and Perturb-seq benchmarks. a, DirAcc (all ground-truth edges) on the six BEELINE datasets for the NT-only, TT-only, and type-matched routing schemes; the TT expert leads on all six datasets, with four reaching DirAcc ≥ 0.83. b, Directional accuracy on the two genome-wide CRISPRi Perturb-seq screens (HCT116 and K562; NT and TT experts) at the default G=200 single window versus genome-wide multi-window coverage; single-window values fall to or below the 0.5 chance level, whereas genome-wide coverage raises accuracy to 0.821 (HCT116) and 0.810 (K562). c, d, Per-condition decomposition for HCT116 (c) and K562 (d): directional accuracy stratified by input scale (G=200 versus full-gene) and edge type (TF→non-TF and TF→TF); the bars labelled “High-conf” denote the retained-edge set at edge_prob > 0.2 (the τ = 0.2 emission threshold), distinct from the optional stricter-precision filter (edge_prob ≥ 0.65) defined in the Methods. On these retained edges accuracy reaches 0.86 (HCT116) and 0.95 (K562), and 0.95/0.96 on the TF→TF subset, whereas shuffled-direction controls return to the chance level (0.5).

#### Rank-deficiency guard for Pearson preselection at low cell-to-gene ratios

Datasets with a low cell-to-gene ratio require a targeted guard to keep correlation-based preselection statistically stable. When the ratio of sampled cells to genes is low, the Pearson correlation matrix tends to be rank-deficient and correlation-based target preselection degenerates into noise-driven ranking, so the candidate gene set must be shrunk by variance. We encountered this with the BEELINE hESC dataset (758 cells, 17,735 genes): its subsampled cell-to-gene ratio of 500/17,735 = 0.028 falls below the preselection stability threshold of 0.04, the Pearson correlation matrix is severely rank-deficient (rank ≤ 500), per-TF top-50 target selection is dominated by noise and effectively random, within-window ground-truth edge coverage collapses, and the genome-wide AUROC is only 0.501 (random level) with an EPR of 1.99×—clearly below the other five datasets (EPR 4.64–5.31). We therefore introduce adaptive variance-based subsampling: when the ratio falls below the threshold, candidates are restricted to the n_keep genes of highest variance (n_keep = n_sub/0.05 = 10,000), compressing the input from 17,735 to 10,000 genes to restore preselection stability. After this fix, hESC’s AUROC rises to 0.718, EPR to 4.19×, and DirAcc to 0.841 (Table 1), bringing its genome-wide results in line with the other datasets (the hESC result in Fig. 3 is the post-fix value). The fix is purely variance-based and uses no ground-truth information; a control benchmark that selects genes by ground-truth connectivity (AUROC 0.736, numerically identical to the single-window hESC value in Fig. 3a) closely matches the fixed result, confirming that the earlier discrepancy reflected preselection rather than the inference model itself. The rank-deficiency guard is therefore a necessary engineering component of window-based preselection at low cell-to-gene ratios, ensuring stable genome-wide coverage under extreme high-dimensional inputs.

#### Design ablations

Systematic ablations confirm that every module of the model contributes an observable performance gain rather than being set heuristically. We first examine the complementarity of the two input feature sets by comparing the P+D dual-channel fusion against single-channel inputs: joint modeling of P and D outperforms the single-channel setting with D features alone (Fig. 5a). Comparing the separated edge/direction training strategy against a joint-training baseline, task-separated training raises edge-prediction AUROC from 0.564 to0.679 (Fig. 5b). Increasing the training corpus from 10k to 20k SCMs causes only a small 2.2% AUROC drop; adding Spearman correlation as a third input channel degrades performance by 7.2%; and superimposing extra noise with σ=0.05 causes a large 24.1% AUROC decline (Fig. 5c). These results indicate that the P+D dual-channel configuration of the precision and distance correlation matrices is sufficient for the tasks evaluated here, and redundant statistics or a larger training set add noise rather than signal. Against the DAGMA constrained training paradigm as a reference baseline, dual-expert routing achieves a DirAcc of 0.910, essentially on par with DAGMA’s 0.905, but with the practical advantage that inference requires no per-dataset optimization (Fig. 5d). A parameter scan over window size G shows AUROC first rising then falling and peaking at G=200 (Fig. 5e). Together, these ablations demonstrate that each design choice of BoYueGRN is empirically justified by an observable performance gain.

**Fig. 5.**
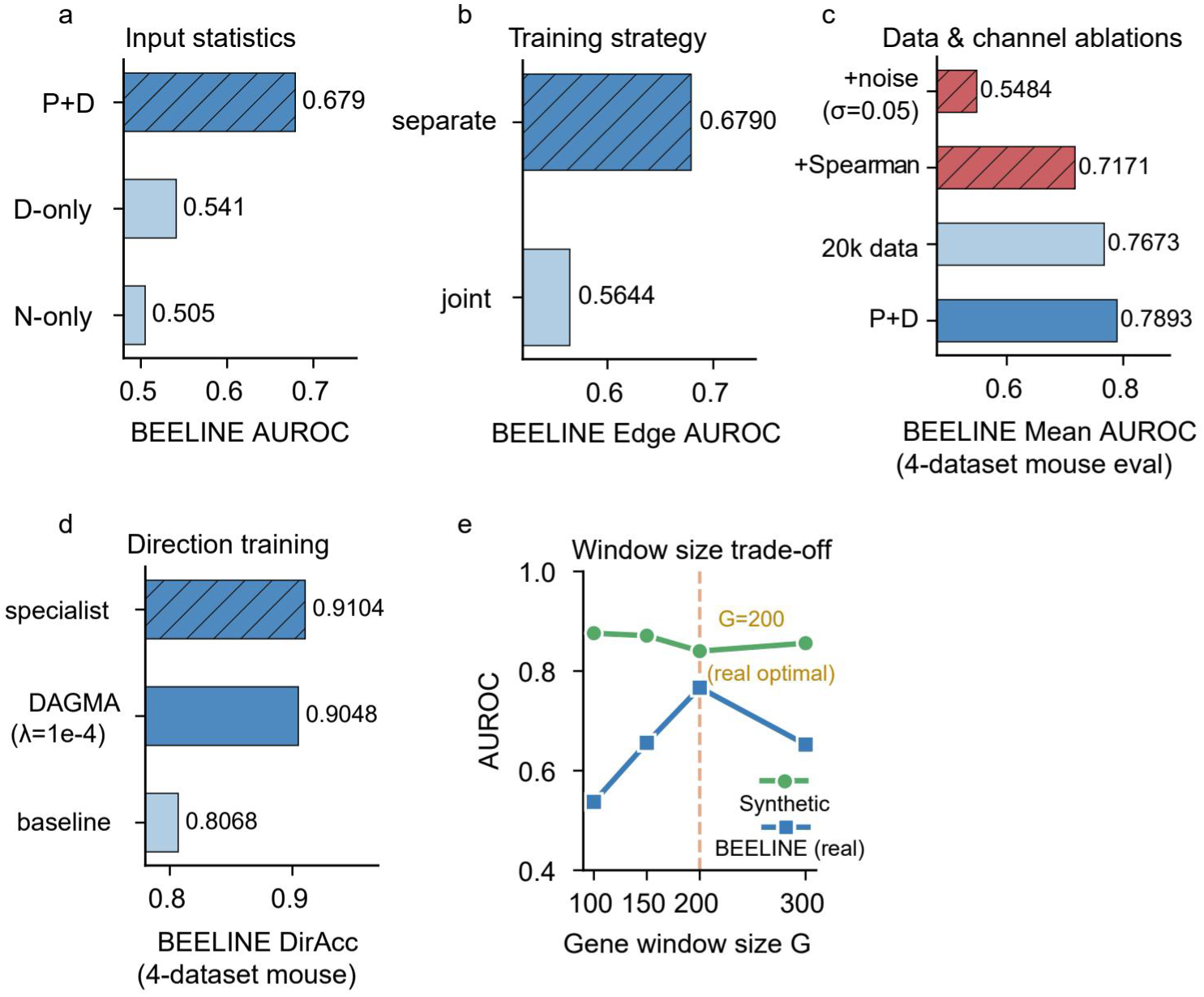
Design ablations. a, Input-statistic ablation (BEELINE AUROC): P+D dual-channel input (0.679) versus D-only (0.541) and N-only (0.505) single-channel inputs. b, Training-strategy ablation (BEELINE edge AUROC): task-separated (0.679) versus joint (0.564) training. c, Data and channel ablations (BEELINE mean AUROC, four mouse datasets): P+D baseline (0.789), 20k-SCM training corpus (0.767), added Spearman-correlation channel (0.717), and superimposed noise with σ=0.05 (0.548). d, Direction-training ablation (BEELINE DirAcc, four mouse datasets): dual-expert specialist (0.910) versus the acyclicity-constrained DAGMA baseline (λ=1e-4; 0.905) and an unspecialized baseline (0.807), with no per-dataset optimization at inference. e, Window-size trade-off: AUROC on synthetic data is flat across G, whereas BEELINE (real) AUROC peaks at G=200.

### Expanded gene selection recovers disease-relevant pathways missed by conventional selection

Because the scale of the input gene set directly constrains biological discovery, we developed a DEG-Expanded strategy that replaces conventional HVG selection and validated it on a multi-stage non-alcoholic fatty liver disease (NAFLD; renamed metabolic dysfunction-associated steatotic liver disease, MASLD, by the 2023 Delphi consensus, with NASH correspondingly MASH) dataset[39]. BoYueGRN trains and infers on fixed-size (G=200) gene subsets and achieves genome-wide coverage through sliding windows, but conventional highly variable gene (HVG) selection typically retains only a few hundred genes (for example 350), a cap that can constrain downstream biological mining. To break this constraint, the DEG-Expanded strategy constructs the input gene set from differentially expressed genes (DEGs). We validate it on a NAFLD cohort (GSE202379, 99,809 cells × 30,908 genes; five stages: Healthy, NAFLD, NASH, Cirrhosis, End-stage). NAFLD is the most common chronic liver disease worldwide and can progress from simple steatosis through steatohepatitis (NASH) to cirrhosis and hepatocellular carcinoma. To isolate the effect of gene coverage on discovery, we merge all cell types, run GO enrichment[40, 41] on the expanded input gene set, and compare term by term against the conventional strategy (DEG-350) (Fig. 6).

**Fig. 6.**
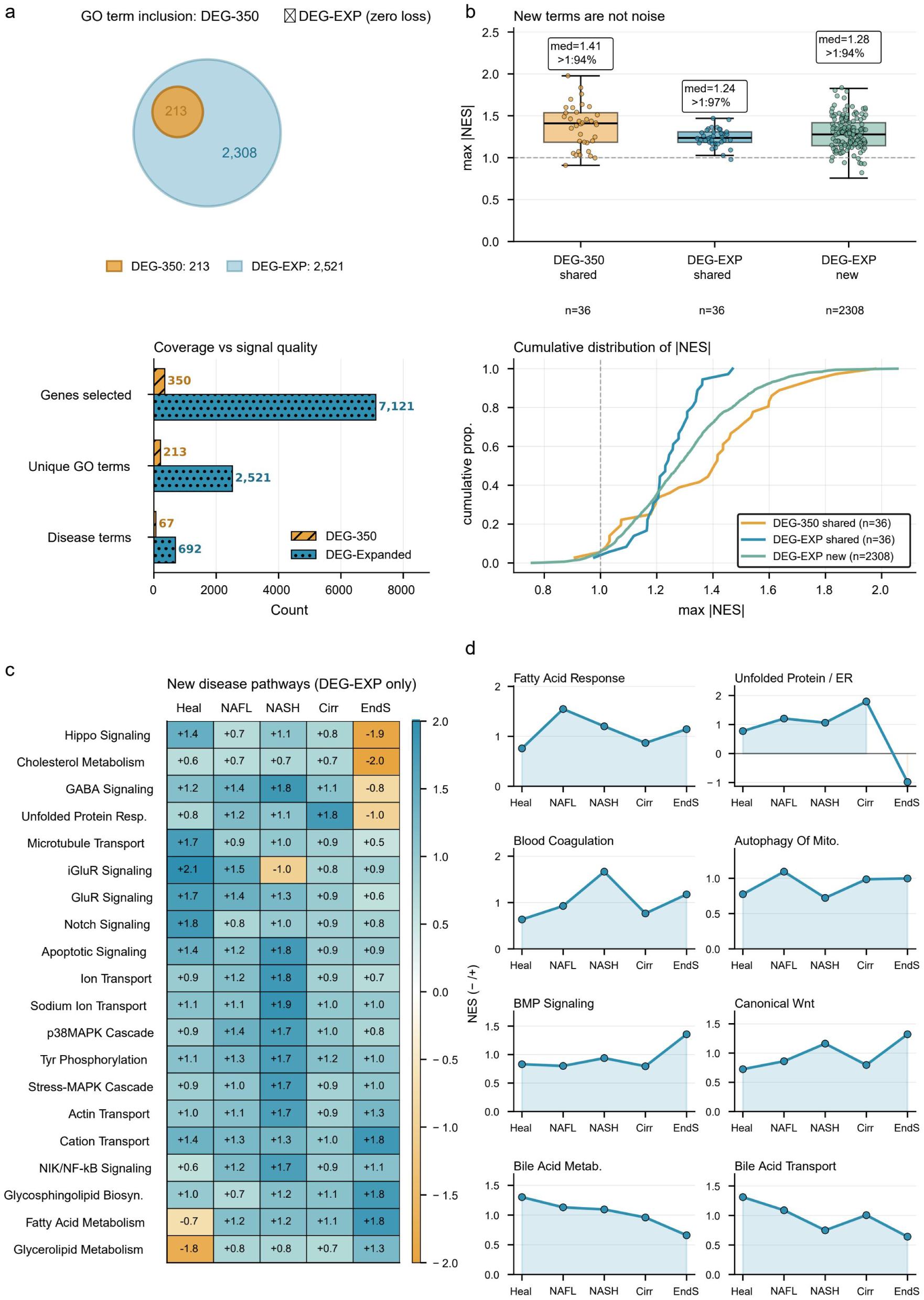
DEG-Expanded gene selection recovers disease-relevant pathways missed by conventional selection (NAFLD). a, GO term inclusion and coverage: all 213 terms detected by the conventional DEG-350 strategy are contained in the 2,521 DEG-Expanded terms (2,308 new), with input genes enlarged from 350 to 7,121 (20.3-fold) and disease-related terms from 67 to 692 (10.3-fold). b, New terms are not noise: max|NES| distributions for DEG-350 shared terms (median 1.41; 94% above |NES|=1), DEG-Expanded shared terms (median 1.24; 97%), and DEG-Expanded new terms (median 1.28; 94%), with cumulative distributions of the three groups. c, Twenty disease pathways detected only under DEG-Expanded form a stage-resolved pathological cascade across Healthy, NAFLD, NASH, Cirrhosis, and End-stage (including Hippo signaling decline, GABA signaling, unfolded protein response, and end-stage glycosphingolipid biosynthesis). d, Cross-stage NES trajectories of eight representative pathways (fatty acid response, unfolded protein/ER response, blood coagulation, autophagy of mitochondria, BMP signaling, canonical Wnt, bile acid metabolism, bile acid transport), showing temporally ordered dominance and pathway off-switching (ER response reversing from activation to suppression at the End-stage).

Expansion broadens coverage without sacrificing specificity (Fig. 6a). All 213 GO terms detected by the conventional strategy (DEG-350) are fully contained in the 2,521 terms of DEG-Expanded, with the input gene number enlarged 20.3-fold and the number of disease-related terms enlarged 10.3-fold. The 2,308 newly added terms carry genuine enrichment signal: their median max|NES| is 1.28, 94% exceed the empirical threshold |NES|=1, and the distribution is comparable to that of shared terms and strongly divergent from random. This coverage expansion translates directly into discoveries inaccessible to the conventional strategy (Fig. 6b). Twenty disease pathways entirely absent under the conventional strategy form a stage-resolved pathological cascade (Fig. 6c): hepatocyte neural/developmental features are lost in the Healthy stage and glutamate receptor signaling declines with progression; regenerative attempts emerge in NASH with co-activation of G2/M transition, liver development, and GABA signaling; genomic instability accompanies immune attack in Cirrhosis, involving DNA/mitotic recombination, T-cell-mediated cytotoxicity, and catecholamine metabolism; and precancerous features of hepatocellular carcinoma (HCC) transformation appear at the End-stage with epigenetic remodeling, including base excision repair, ganglioside metabolism, regulation of histone modification, and telomere-maintenance suppression. Ganglioside metabolism, strongly activated at the End-stage with little precedent in the NAFLD literature, is a candidate HCC marker. Cross-stage trajectories of eight representative pathways further validate the temporal consistency of the outputs (Fig. 6d): lipid response (NAFL), coagulation (NASH), ER stress (Cirrhosis, reversed at End-stage), and BMP/Wnt signaling (End-stage) dominate in turn, consistent with classical pathology, and ER stress shifting from activation to suppression shows that the method captures pathway off-switching, not only activation. These results indicate that conventional gene-selection strategies systematically lose disease-mechanism-relevant signals, whereas DEG-Expanded leverages expanded gene coverage to mine reproducible, evidence-backed pathway features.

### BoYueGRN reveals stage-specific regulatory programs in disease progression

Having fixed a concrete analysis protocol through the DEG-Expanded evaluation—DEG-Expanded gene selection, the TT direction expert, and an edge_prob > 0.2 threshold, we apply BoYueGRN to five diseases spanning metabolic (NAFLD)[39], chronic inflammatory (periodontitis)[42], neoplastic (HCC)[43], precancerous (proliferative verrucous leukoplakia, PVL)[44], and neurodegenerative pathologies (Alzheimer’s disease, AD)[45]. Each disease spans multiple stages and cell types (Fig. 7–8, Fig. S4–6). We resolved each dataset by cell type and stage, examining which stage-specific regulatory programs appear in each cell type and which TFs drive them. Directed, genome-wide GRNs are well suited to these questions: edge-count trajectories track program rewiring, and driver-TF reverse lookup identifies the TFs behind each pathway change. Together the five settings provide a broad test of the method’s generality.

**Fig. 7.**
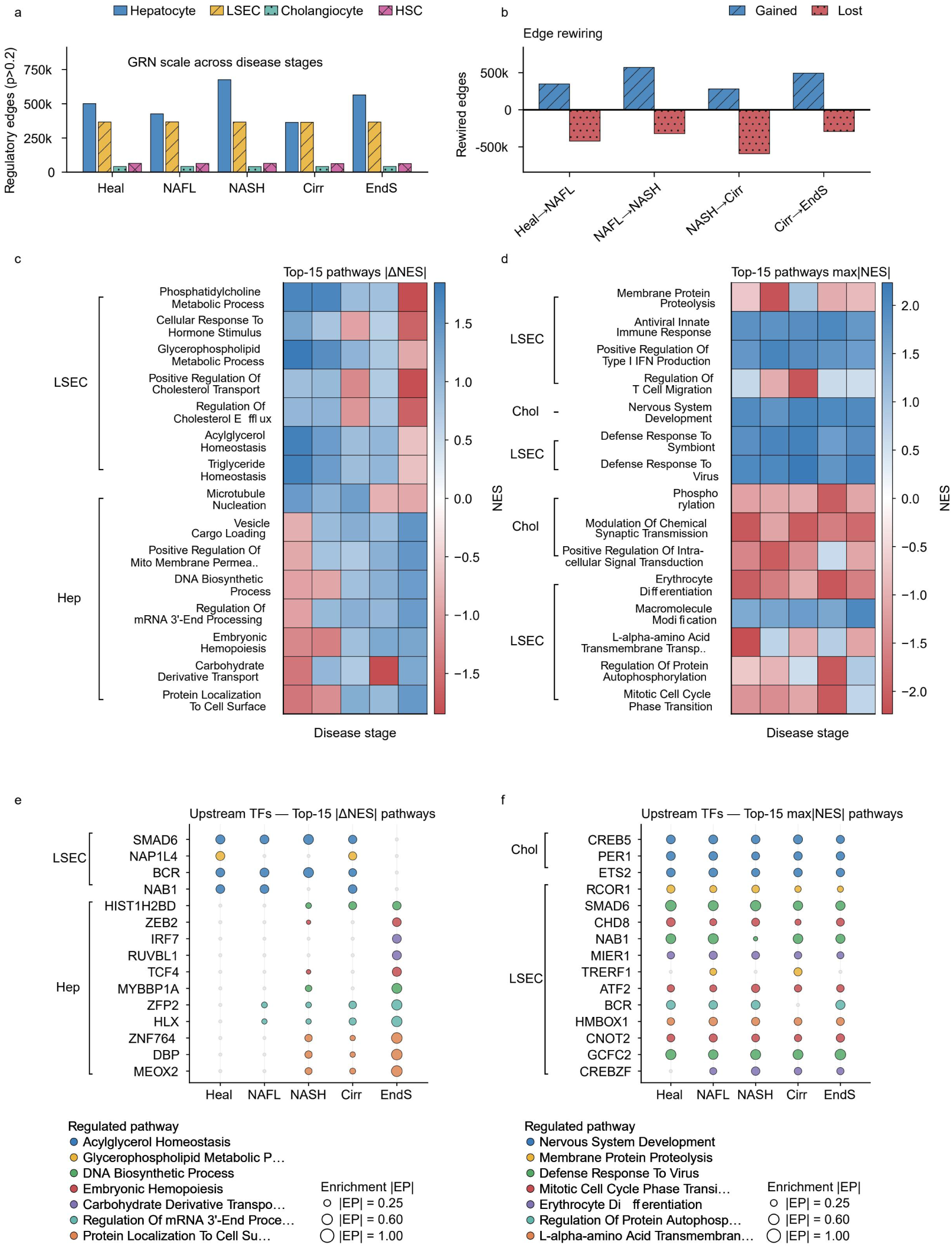
Cell-type-resolved regulatory rewiring across NAFLD progression. a, GRN scale (regulatory edges, p>0.2) across disease stages by cell type: hepatocytes follow a pronounced biphasic expansion (501k edges at Healthy, 676k at NASH, 364k at Cirrhosis, 565k at End-stage), whereas LSECs, cholangiocytes, and hepatic stellate cells remain nearly constant. b, Edge rewiring (gained and lost edges) between consecutive stage transitions: NAFLD-to-NASH is the largest expansion and NASH-to-Cirrhosis the largest contraction. c, d, Top-15 pathways ranked by |ΔNES| (c) and max|NES| (d) across stages for LSECs and hepatocytes, including end-stage collapse of LSEC lipid/cholesterol metabolic programs with maintained antiviral/innate immune activation, and hepatocyte DNA-synthesis and mitochondrial-membrane-permeabilization programs. e, f, Upstream regulators of the top pathways ranked by |ΔNES| (e) and max|NES| (f): the LSEC lipid-pathway collapse is driven by synchronous decline of SMAD6 and NAP1L4, while MEOX2, DBP, and TCF4 establish de novo regulation.

**Fig. 8.**
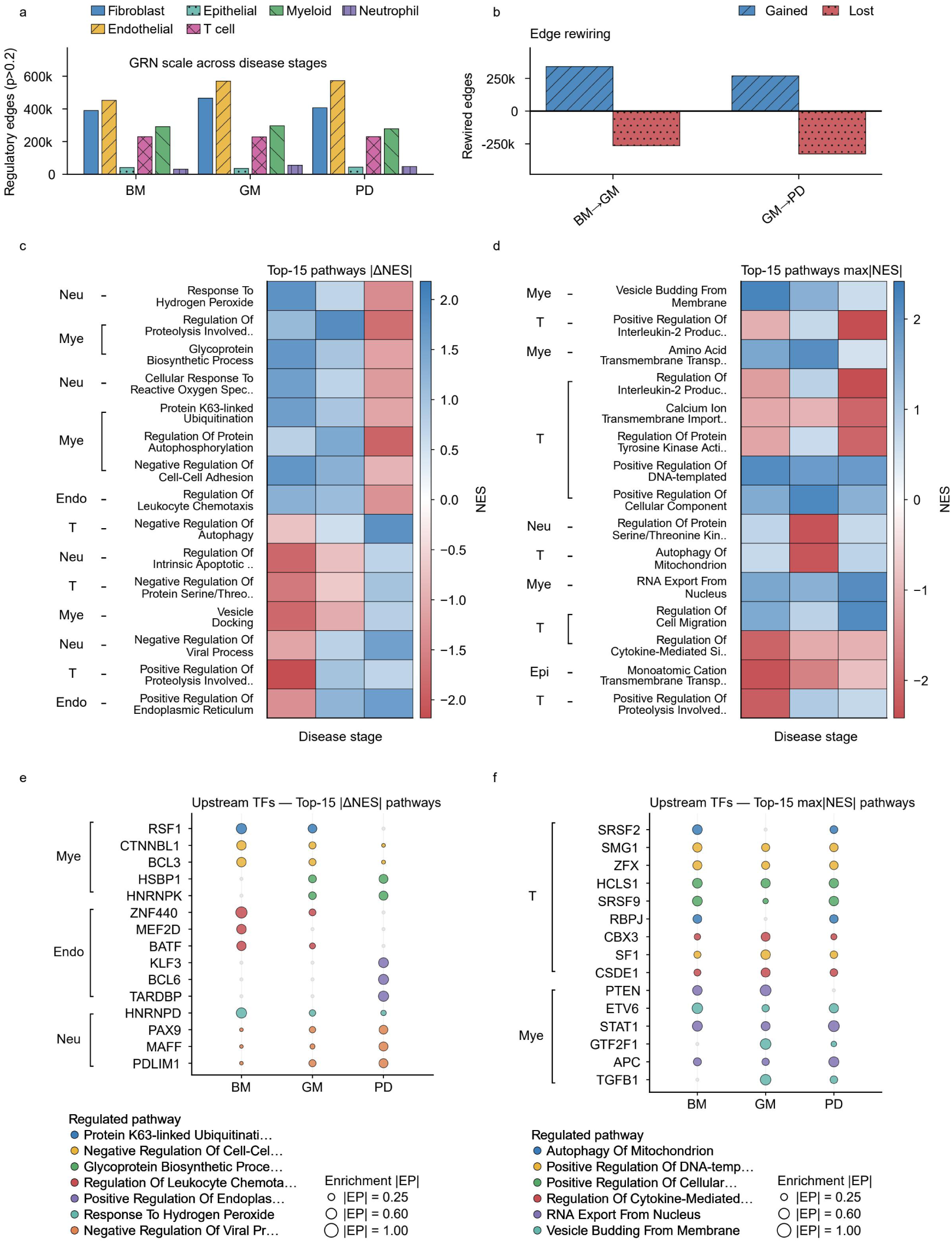
A reversible-to-irreversible regulatory state transition in periodontitis. a, GRN scale across stages (healthy buccal mucosa BM, gingivitis GM, periodontitis PD) for six cell types: fibroblasts show an inverted-U trajectory (390k edges at BM, 466k at GM, 407k at PD), whereas the epithelial GRN remains stable. b, Edge rewiring: BM-to-GM progression nets 75.6k gained edges; GM-to-PD progression nets 58.6k lost edges. c, d, Top-15 pathways ranked by |ΔNES| (c) and max|NES| (d) across stages and cell types; the edges lost at PD are mainly attributable to collagen biosynthesis and tissue repair while inflammatory programs persist. e, f, Upstream regulators of the top pathways ranked by |ΔNES| (e) and max|NES| (f): myeloid cells are dominated by weakening of cell-cycle and protein-ubiquitination programs (including RSF1), T cells are led by the splicing factors SRSF9 and SRSF2, and the strong myeloid signal of TGFB1 (a master fibrosis regulator) corresponds to the fibrosis axis of periodontitis; the complete per-cell-type rankings (including the myeloid innate-immune regulator TLR4) are given in the supplementary driver-TF panels (Supplementary Fig. S7).

#### Biphasic regulatory network rewiring in NAFLD progression

Cell-type-resolved analysis shows that NAFLD regulatory dynamics are driven almost entirely by hepatocytes (Fig. 7a). The hepatocyte GRN size follows a pronounced biphasic expansion trajectory: 501k at Healthy, peaking at 676k at NASH, bottoming at 364k at Cirrhosis, and rebounding to 565k at End-stage, while LSECs, cholangiocytes, and hepatic stellate cells remain nearly constant. Edge rewiring localizes the two turning points to specific stage transitions: NAFLD-to-NASH is the largest expansion period and NASH-to-Cirrhosis the largest contraction (Fig. 7b). Cell-type-resolved pathway and driver analyses uncover three programs invisible in merged analyses (Fig. 7c–f). First, LSECs collapse their lipid/cholesterol metabolism at the End-stage while maintaining strong activation of antiviral/innate immune pathways, consistent with the established scavenger and antigen-presentation roles of liver sinusoidal endothelial cells, with the end-stage timing of the lipid-program collapse being the novel element contributed by the present analysis. Second, hepatocytes simultaneously activate DNA synthesis and mitochondrial membrane permeabilization at the End-stage, complementing the regenerative signals seen at the merged level and corresponding to regenerative failure in end-stage liver disease. Third, the LSEC lipid-pathway collapse is driven by the synchronous decline of related TFs such as SMAD6 and NAP1L4, while MEOX2, DBP, and TCF4 establish *de novo* regulation, mutually corroborating the HCC-transforming precancerous pathways recovered by DEG-Expanded enrichment at the End-stage and consistent with the NASH-to-HCC transition reported in the literature. The network also reproduces the literature consensus patterns of declining Hippo signaling across NAFLD stages and canonical Wnt activation at the End-stage[46], independently supporting the biological fidelity of cell-type-resolved GRNs.

#### A reversible-to-irreversible regulatory state transition in periodontitis

Cell-type-resolved GRN analysis of a public periodontal single-cell dataset (GSE164241, 51,110 cells × 23,341 genes; three stages: healthy buccal mucosa, BM, gingivitis GM, periodontitis PD) reveals a switch from a reversible to an irreversible regulatory state. Periodontitis is among the most common chronic inflammatory diseases of the oral cavity; early gingivitis is fully reversible by plaque removal, whereas progression to periodontitis entails irreversible alveolar bone resorption and attachment loss. Fibroblasts, the core effectors of periodontal tissue repair and ECM homeostasis, show an inverted U GRN size trajectory, expanding from 390k edges at the healthy stage to a peak of 466k at GM, then falling back to 407k at PD, whereas the epithelial GRN size remains stable (Fig. 8a). Edge-rewiring analysis reveals two qualitatively different regimes: BM to GM progression nets 75.6k edges with co-activation of inflammatory response and ECM organization programs, corresponding to a reversible inflammatory-repair response to plaque stimulation, whereas GM to PD progression nets a loss of 58.6k edges, with the lost edges mainly attributable to collagen biosynthesis and tissue repair while inflammatory programs persist, suggesting that the irreversibility of PD tissue destruction stems not from enhanced inflammation but from active silencing of repair programs (Fig. 8b–d). Driver-TF reverse lookup (Fig. 8e–f) shows that periodontitis drivers are distributed across multiple non-epithelial compartments rather than concentrated in a single cell type. Ranked by regulatory-change magnitude, myeloid cells are dominated by weakening of cell-cycle and protein ubiquitination programs (RSF1 and other myeloid regulators) and endothelial cells by rearrangement of autophagy and glycan-metabolism programs (ZNF440 and other endothelial regulators). Ranked by signal strength, the top-ranked regulators in Fig. 8f include GTF2F1, ETV6, TGFB1, and the T-cell splicing factors SRSF9 and SRSF2. The strong myeloid signal of TGFB1 (a master fibrosis regulator), together with the innate-immune programs prominent in myeloid cells, corresponds to the fibrosis and inflammation axes of periodontitis (Fig. 8e–f). At the network level, the inverted-U trajectory identifies gingivitis as a reversible window: intervention while repair programs remain active can restore tissue homeostasis, whereas once repair programs are silenced, damage accumulates and cannot self-reverse, providing a GRN-level account consistent with the established clinical reversibility of gingivitis, a hypothesis-generating observation rather than new clinical evidence.

#### Extended analyses of neoplastic, precancerous, and neurodegenerative settings

Extending the same analysis to three additional pathologies confirms that BoYueGRN consistently recovers pathology-coherent programs across diverse disease modes. HCC is the natural extension of the NAFLD case: the precancerous pathways and MYC expansion detected at the NAFLD End-stage are fully expressed in HCC, and the two cases form an internally consistent progression chain. In HCC (Normal and Tumor), the hepatocyte GRN size expands 21-fold in an almost purely additive manner (Fig. S4a, b), and the pathway heatmaps reveal a metabolic-to-proliferative switch, with DNA-replication and cell-cycle programs activated and hepatocyte-specific metabolic programs suppressed (Fig. S4c, d). In driver-TF reverse lookup, the core drivers NCOA6, DNAJC2, and SNRPB, independently returned by the change-magnitude (Fig. S4e) and signal-strength (Fig. S4f) rankings, pointing to metabolic reprogramming, ribosomal stress, and splicing dysregulation. PVL, an invasive oral precancerous lesion prone to recurrence and malignant transformation, shows a highly cell-type-heterogeneous response across its mild/moderate/severe stages (with healthy gingival BM reused from the periodontitis dataset as control, so cross-study batch and site differences may contribute to the reported expansion ratios): the T-cell GRN expands 5.6-fold and the epithelial GRN 2.3-fold, with core perturbations in the microenvironmental compartment, and driver-TF reverse lookup independently recovers IRF6 (Fig. S5)[47]. In AD (Control and AD; Fig. S6), network expansion is mild; at the pathway level, the most prominent change in astrocytes is enhancement of protein-complex assembly and protein-transport programs, accompanied by weakening of MAPK/ERK cascade signaling, while oxidative phosphorylation, insulin-receptor signaling, and synaptic-organization terms appear but do not reach significance. Together, the consistent recovery of pathology-coherent programs across neoplastic, precancerous, and neurodegenerative settings supports the general applicability of the method.

## Discussion

BoYueGRN reframes directed GRN inference as a train-once, cross-dataset inference paradigm, replacing the per-dataset optimization that has long constrained the field. Trained entirely on synthetic structural causal models, the model reconstructs genome-scale directed regulatory networks from real scRNA-seq data in a single forward pass, simultaneously delivering the prediction accuracy, gene coverage, and reliability of regulatory direction that existing tools struggle to combine. We validated its performance on the BEELINE benchmark and two Perturb-seq screens, and, applying the same pipeline to five disease datasets, the model not only reproduced reported pathological features but also generated experimentally testable regulatory hypotheses, demonstrating both reliable predictive ability and mechanistic discovery value.

To address the high-dimensionality, noise, and directional constraints inherent to scRNA-seq GRN inference, we adopt an amortized causal discovery design. GENIE3, GRNBoost2, graphical lasso, and NOTEARS all refit models per dataset, which is slow and unstable when p≫n; an amortized model trains once and then infers at near-zero marginal cost. This lineage extends from the time-series-oriented ACD and the i.i.d.-setting AVICI to the CDFM foundation model[29, 30, 32], yet none targets GRN inference, and even the closest predecessor, AVICI, has been validated only on synthetic and semi-synthetic data. What enables the transfer is the alignment of the synthetic training distribution with real GRNs along three axes: (1) scale-free hub topology from preferential attachment; (2) an additive-noise mechanism that separates injection noise from final noise to render directions identifiable; and (3) log1p-transformed, sparse, non-negative expression statistics. Whereas simulators such as SERGIO generate gold-standard benchmarks for GRN methods[31], BoYueGRN instead uses synthetic causal models directly as training material and shows that effective mechanistic alignment between synthetic data and real single-cell transcriptomes enables zero-shot transfer inference to real datasets.

Four design decisions jointly underpin the model’s performance. First, fusing the Ledoit–Wolf shrunk precision matrix with the distance correlation matrix jointly captures linear and nonlinear regulation, outperforming the distance-correlation-only channel. Second, training edge existence and direction separately, because the two tasks demand conflicting features, raises edge-prediction AUROC from 0.564 to 0.679, and TT/NT dual-expert routing achieves a DirAcc of 0.910, matching the acyclicity-constrained DAGMA (0.905) while requiring no optimization at inference[48]. Third, TF-centric sliding windows with asymmetric fusion extend the fixed 200-gene model to 99.4–100% genome-wide coverage without loss of accuracy. Fourth, replacing conventional HVG selection with DEG-Expanded enlarges the NAFLD input 20-fold while preserving every GO term detected by the conventional strategy, shifting analysis from genome-wide scanning toward disease-focused reconstruction.

Applying the framework to GRN-dynamics reconstruction across five diseases spanning neoplastic, precancerous, and neurodegenerative settings recovered pathology-coherent regulatory programs and generated testable hypotheses in every case, with most of these signals recovered only at cell-type resolution rather than in merged analyses. Because inference requires no retraining or case-specific tuning, the same zero-shot pipeline applies directly to any disease with scRNA-seq data, making BoYueGRN broadly accessible for hypothesis generation and mechanistic discovery. Furthermore, we provide the top 15 aggregated results across cell types. Users may select cell types of interest to perform in-depth mining of cell-specific regulatory features. For instance, analyses of PVL T cells reveal markedly enhanced regulatory activity of IRF6, a well-validated tumor suppressor in squamous carcinoma, from mild to moderate lesion stages, with its regulated target genes expanding from 88 to 470, consistent with tumor-suppressive programs being engaged rather than lost during early lesion progression. As illustrated by this, a single set of GRN inferences can support both broad-scale summaries and cell-type-focused analyses.

BoYueGRN can deliver reliable GRN-inferred outputs for downstream tasks. The most direct extension is *in silico* perturbation simulation on BoYueGRN’s directed regulatory networks, predicting transcriptome-cascade effects of TF knockout or overexpression to computationally prioritize CRISPR functional screens. Integrating the framework with multi-omics signals such as ATAC-seq motif analysis and spatial-transcriptomic neighborhood information could further improve tissue-specific regulatory inference. At the mechanistic level, the multi-cell-type, stage-resolved directed GRNs produced here provide a new basis for dissecting intercellular regulatory crosstalk—resolving transcriptional signal transmission between cells, distinguishing cell-autonomous from non-cell-autonomous drivers, and tracing the origin nodes of regulatory dysregulation across disease stages. At the translational level, GRN-dynamics features such as network expansion ratio are candidates for prognostic modeling once patient-level outcome data become available; the present analyses are hypothesis-generating and carry no clinical validation.

It is worth noting that although we have implemented corresponding countermeasures, some inherent challenges of GRN inference cannot be fully eliminated. The gold-standard annotations of the BEELINE benchmark, derived from ChIP-seq, suffer incomplete coverage: many so-called negative samples merely reflect absent evidence rather than proven absence of regulation, lowering the attainable AUROC ceiling[25]. We therefore use both AUROC and EPR in all evaluations, and interventional benchmarks in the style of CausalBench would provide a stronger basis for causal claims and are a natural next step. In addition, DEG calls are computed per cell with Wilcoxon tests and do not model sample-level nesting, so significance may be inflated; DEG selection is therefore used as a gene-coverage heuristic rather than an inferential claim, and pseudobulk or mixed-model sensitivity analyses are a worthwhile refinement. In addition, cell types with few cells and few differential genes are subject to limited statistical power, and the TF number and window-rotation mechanism of the sliding-window framework introduce secondary biases. Further, the CRISPRi-based Perturb-seq ground truth is itself conservative: both screens use knockdown rather than knockout, the median knockdown efficiency in HCT116 is approximately 75.4%, weaker regulatory effects are hard to detect, and measured directional accuracy therefore likely underestimates BoYueGRN’s true performance. Furthermore, the cell-type-specific GRNs of the five disease datasets are currently validated mainly through indirect consistency with the published literature, and some newly generated hypotheses await confirmation in independent cohorts or wet-lab perturbations, a limitation imposed by the high cost of in vivo cell-type-specific perturbation experiments and shared by single-cell research broadly rather than specific to this method.

In summary, BoYueGRN demonstrates that amortized training on synthetic data can learn causal features that transfer zero-shot to real scRNA-seq, even without epigenetic priors or perturbation information, reconstructing biologically valid directed regulatory networks from steady-state transcriptomes. This work shifts GRN inference from a per-dataset optimization paradigm to a train-once-reuse paradigm and provides a unified framework for multi-stage, multi-cell-type disease regulatory dynamics. As single-cell multi-omics data and large-scale perturbation screens continue to accumulate, this paradigm—combining prior-free direction inference, genome-wide coverage, and disease-network dynamic phenotype analysis—should prove valuable for disease-mechanism elucidation, therapeutic target discovery, and precision medicine.

## Methods

### 1. Framework overview

BoYueGRN is an amortized GRN inference framework trained once on synthetic SCMs and thereafter producing a directed, genome-wide GRN for any new real dataset in a single forward pass. The workflow proceeds in three stages. First, synthetic data generation constructs 10,000 expression datasets under controllable rules with fully known graph structure, edge weights, mechanism types, and noise magnitudes, providing complete supervision for both the edge-existence and direction tasks. Second, model training serves the edge-existence and direction tasks with independently trained encoders, the direction task further split into dual experts by target-gene type. Third, zero-shot inference slides fixed G=200-gene windows TF-centrically across the genome, one forward pass per window, fusing edge probability by maximum and direction score by mean, and emitting directed edges at threshold τ=0.2. The upfront cost of the amortized paradigm buys nearly free inference at deployment—the computational prerequisite for large-scale GRN-dynamics analysis across cell types, stages, and diseases.

### 2. Data sources and preprocessing

#### Data sources

Model training uses only synthetic data (Section 3); all real data are reserved for zero-shot evaluation and biological applications. The first class is the BEELINE benchmark, comprising six scRNA-seq datasets (mDC, mHSC-E, mHSC-GM, mHSC-L, hESC, hHep) with the ChIP-seq-derived directed networks released with the benchmark as ground truth, for fair comparison against eight existing methods (GENIE3, GRNBoost2, Correlation, MutualInfo, PPCOR, GLasso, dCor, and a random baseline)[25]. The second class is the Perturb-seq benchmark, comprising the X-Atlas/Orion genome-wide CRISPR interference (CRISPRi) Perturb-seq screen in HCT116 (colon cancer cell line, 38,606 genes, about 3.4 million cells)[37] and the genome-wide CRISPRi screen of Replogle et al. in K562 (myeloid leukemia cell line, 8,248 genes, 11,258 cells)[38], both providing causal ground truth for regulatory direction through TF knockdown with transcriptomic-response measurement. The third class is disease cases: NAFLD (GSE202379)[39], hepatocellular carcinoma (HCC, GSE149614)[43], periodontitis (GSE164241) [42], Alzheimer’s disease (GSE157827)[45], and proliferative verrucous leukoplakia (GSE196296) [44].

#### Preprocessing

Each expression matrix is z-scored per gene before feature computation (Section 4). Cell filtering proceeds in two steps: Perturb-seq cells are grouped by guide type with at least 50 cells per perturbed TF, and disease-case cells are grouped by cell-type annotation with at least 100 cells per (cell type × stage) combination (the “valid cell type” criterion). At most 500 cells per window are used for input-feature computation (MAX_CELLS=500), with downsampling beyond that, to control the cost and numerical stability of precision-matrix estimation. For Perturb-seq ground truth, HCT116 uses a Welch t test to define target genes significantly differentially expressed after TF knockdown (p < 0.01, |log2FC| > 0.25; 1,320 perturbed TFs); because CRISPRi mediates transcriptional knockdown rather than complete knockout, the median knockdown efficiency in HCT116 is approximately 75.4%[37], and these ground-truth edges may underestimate true regulatory effects. K562 follows the authors’ released Truth-seq calls (|z| ≥ 2.0) to define regulatory edges (663 TFs covered by windows).

### 3. Synthetic data generation with structural causal models

Synthetic data are generated by structural causal models under controllable rules (10,000 independent datasets; each G=200 genes × C=1,000 cells), providing the complete ground truth that real scRNA-seq lacks, since only a small fraction of gene pairs can be validated by ChIP-seq or Perturb-seq. **Graph topology.** The first five nodes form a seed causal chain (each node has exactly one incoming edge from an earlier-indexed node); subsequent nodes sample parents among earlier-indexed nodes with probability proportional to (in-degree + 1), with an expected parent count of about 5. Constraining parents to earlier-indexed nodes guarantees acyclicity by construction, and preferential attachment[34] produces a scale-free hub structure mimicking the real-GRN characteristic in which a few hub TFs regulate many targets. Edge weights W ∼ Uniform(−1.5, 2.0), mixing activation (about 57%) and repression (about 43%). **Structural equations.** Each node’s expression is generated in topological order through additive noise structural equations (ANM)[27], with about 50% linear and about 50% GELU mechanisms.

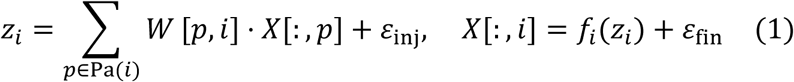

The magnitude gap between the injection noise ε_inf_ ∼ *<u>N</u>*(0, 0.3^2^) and the final noise ε_fin_ ∼ *N*(0, 2.0^2^) satisfies the directional identifiability condition of additive noise models: independence holds in the parent-to-child direction but not the reverse, providing an identifiable signal for direction learning, further reinforced by the 50% nonlinear mechanisms. After log1p transformation of the non-negative expression, synthetic expression acquires the non-negative, sparse, approximately log-normal statistics of scRNA-seq (about 37% zeros), aligning the synthetic distribution with real single-cell data. The 10,000 datasets are split 80/10/10 (train/validation/test) under fixed random seeds, consistent across training seeds.

### 4. Input features

Two complementary gene-pair features are computed from the standardized expression of the G genes in a window: the Ledoit–Wolf shrunk precision matrix P (linear conditional dependence) and the distance correlation matrix D (nonlinear dependence). P is the inverse of the Ledoit–Wolf shrunk covariance estimate; P[i,j] reflects partial correlation after removing confounding by the remaining G−2 genes, capturing direct regulation rather than indirect co-expression, and shrinkage guarantees invertibility and numerical stability when G approaches the sample size. D is defined by the covariance of double-centered distances (dCov²).

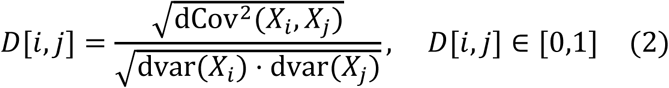

D captures nonlinear/non-monotonic dependence inaccessible to linear partial correlation, making the two channels complementary. The model input is the concatenation cat([P, D]) of shape (1, G, 2G), linearly projected to d_model=512 before entering the graph Transformer encoder.

### 5. Model architecture

BoYueGRN uses three independently trained encoders and two independent prediction heads. The three encoders share one architecture—an 8-layer graph Transformer[49] (GraphTransformerEncoder, G=200, d_model=512, n_heads=8, n_layers=8, dropout=0.1, with edge-biased attention and stochastic depth), each gene a node. The edge encoder is trained from scratch, while the two direction encoders are warm-started from its weights and fine-tuned independently (with no final parameter sharing), because the edge and direction heads require conflicting features (distinguishing edge/no-edge versus distinguishing i→j from j→i). The edge-existence head (EdgeHead; about 25.4 million parameters in total) scores gene pairs with a symmetric dot product superimposed on pairwise statistics.

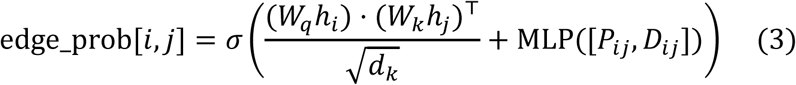

The direction head (AsymmetricDirHead; about 1.05 million parameters) scores direction with asymmetric source/target projections.

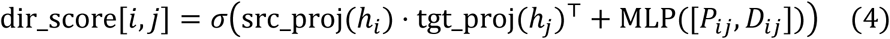

Because the two projections differ, the ordered-pair scores dir_score[*i*, *j*] and dir_score[*j*, *i*] are decoupled by construction, structurally requiring features that distinguish the regulator from the target role. The direction head is further trained as dual experts by target-gene type: the TT expert only on TF-to-TF ground-truth edges and the NT expert only on TF-to-non-TF edges, with inference routed by target-gene type (TF targets to the TT expert, non-TF targets to the NT expert). Because the TT expert outperforms the NT expert on both edge types in genome-wide HCT116 validation, the benchmark and disease-case analyses reported here uniformly use the TT expert.

### 6. Training procedure

All models are trained only on synthetic data. **Edge model.** 20,000 steps, batch size 32, Adam optimizer[50] (lr=2e-4, weight_decay=0.01, linear warmup of 2,000 steps), gradient clipping 1.0; the loss is positive-example-weighted binary cross-entropy on off-diagonal gene pairs with label smoothing,

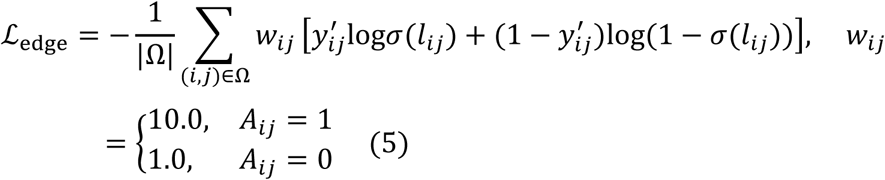

where *y_ij_*’= 0.9*A_ij_* + 0.1(1 − *A_ij_*) (label_smoothing=0.1, shifting each target 0.1 toward the opposite class) and *A_ij_* is the ground-truth adjacency; positive weighting (pos_weight=10.0) mitigates the extreme class imbalance in which positive edges account for only about 0.1%. Four random seeds (0–3) are trained independently and their predictions averaged at inference. **Direction model.** 20,000 steps, Adam (lr=1e-4, weight_decay=0.01, linear warmup of 1,000 steps), label-smoothed binary cross-entropy (smoothing=0.1) computed only on ground-truth edges of the corresponding type,

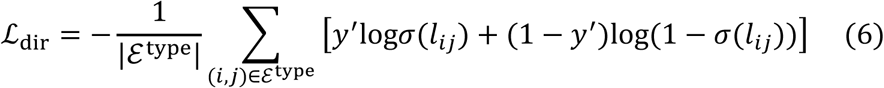

where ℰ^type^ is the TT (TF-to-TF) or NT (TF-to-non-TF) ground-truth edge set and the direction label is the edge direction (1 for i→j, 0 for j→i); the NT expert is a four-seed ensemble and the TT expert uses the best single seed. No real scRNA-seq data are seen during training; zero-shot generalization relies on alignment of the synthetic distribution with real single-cell statistics (Section 3).

### 7. Zero-shot inference and genome-wide coverage

For gene sets larger than 200, fixed TF-centric windows slide across the gene set to achieve genome-wide coverage. Each window contains 5 TFs (TF_PER_WINDOW=5) and 195 target genes, with targets preselected by Pearson correlation (top-50 candidates per TF on a subsample of n_sub = min(500, n_cells) cells) and rotated across windows so every gene is sufficiently covered; each window performs one independent forward pass (about 1 ms). Cross-window fusion follows a complementary rule: edge probability is fused by the maximum across windows (a recall-oriented strategy retaining strong evidence from any window), and direction score by the mean across windows (a multi-window consensus suppressing single-window noise).

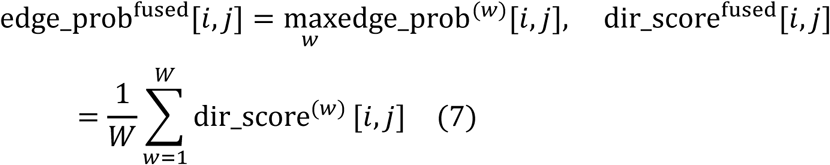

Directed edges are emitted at threshold τ=0.2 (an empirical value balancing precision/recall and consistent with the low-confidence bin boundary of the calibration curve). **Rank-deficiency guard.** The statistical stability of preselection depends on the cell-to-gene ratio n_sub/n_genes (n_sub = min(500, n_cells)); when the ratio falls below 0.04, the Pearson correlation matrix is rank-deficient (rank ≤ n_sub) and top-50 target selection is noise-dominated, so candidates are restricted to the n_keep = n_sub/0.05 genes of highest variance (e.g., for hESC with ratio 500/17,735 ≈ 0.028 below 0.04, n_keep = 10,000). The guard is based only on gene variance and uses no ground-truth information.

### 8. Benchmark evaluation

#### BEELINE zero-shot benchmark

The six datasets are compared against the same seven methods plus a random baseline on identical inputs. BEELINE additionally provides a seventh dataset (mESC), which we exclude because it is the only time-course entry in the benchmark: its 421 cells pool five differentiation time points (0-72 h), violating the steady-state, exchangeable-cells assumption on which the input statistics P and D rest, whereas the other six datasets are single-condition snapshots. The window-level benchmark uses a G=200 single window (top-50 TFs + the 150 target genes of highest expression variance, padded with the highest-variance genes when fewer than 200), ensuring fair comparison with baselines; the genome-wide benchmark covers all genes with the Section 7 sliding windows. Metrics are AUROC, EPR, ECR, and DirAcc (defined in Section 11). **Perturb-seq direction validation.** Because ChIP-seq ground truth reflects physical binding rather than causal regulation, direction inference is further validated on two human Perturb-seq datasets (HCT116 and K562; ground-truth construction in Section 2). Directional accuracy is computed only on predicted edges (edge_prob > 0.2); shuffled controls (randomly permuted ground-truth directions) test for systematic bias in direction judgment. **Confidence calibration.** Predicted edges are binned by edge_prob into low/mid/high bins (bin center 0.10/0.50/0.90), and the empirical DirAcc per bin is computed to test consistency between model probability and independently validated accuracy; an optional high-confidence filter (edge_prob ≥ 0.65) can further restrict edges when stricter precision is desired, although the analyses reported here operate on the τ=0.2 emission threshold (Section 7).

### 9. Disease-case analyses and the DEG-Expanded strategy

The fixed window scale G=200 also determines the gene-selection strategy, because gene selection dictates which genes enter the sliding windows and hence the network’s coverage. Conventional HVG selection restricts the input to a few hundred genes (for example 350), a cap that limits the discovery range by excluding many disease-relevant genes (mid- and low-expression markers, cell-type-specific TFs). We therefore propose the DEG-Expanded strategy, constructing the input from statistically significant differentially expressed genes to lift the cap. **Unified pipeline.** All five cases share one pipeline. DEGs are first selected by Wilcoxon rank-sum test (stage vs baseline, BH-FDR < 0.05, |log2FC| > 0.5), with the union across stages taken and no truncation (e.g., 7,121 genes for NAFLD). TFs are then filtered from the DEG set using a species TF list (the BEELINE human TF catalog compiled from RegNetwork and TRRUST, which includes curated non-classical regulators such as signaling molecules, receptors, and splicing factors alongside DNA-binding transcription factors); when the cap is exceeded, the top max_tfs=150 are taken in descending |log2FC| order. Direction inference uses the TT expert; sliding windows, preselection, fusion, and the edge threshold follow Section 7 (edge_prob > 0.2). **Cell type × stage resolution.** Each dataset is split by cell-type annotation and stage, and one GRN is inferred per (cell type × stage) combination (≥ 100 cells per group), preventing merged analyses from canceling cell-type-specific regulatory signals; subsequent pathway and driver analyses operate at this resolution.

### 10. GO enrichment and pathway interpretation

For each (cell type × stage) GRN, a prerank gene list is built using TF-to-target edge probability as the gene-importance score, and prerank GSEA (gseapy implementation) is run against GO biological-process terms, reporting each term’s NES, nominal p-value, and FDR q-value. **Filtering rules.** Enrichments dominated by ribosomal gene families (RPL/RPS/MRPL/MRPS; housekeeping-gene contamination) and overly broad generic transcription terms (e.g., GO:0006355) are removed; NOM p < 0.05 is primary and p < 0.10 secondary. **Pathway selection and ranking.** Pathways are ranked by |ΔNES| (late−early; the most drastic changes) and by max|NES| (strongest cross-stage signal), taking the top 15 of each, deduplicating across cell types, and retaining the best cell type. **Driver-TF reverse lookup.** For each selected pathway, the edge_prob of every TF over all target genes in the pathway is aggregated; e2 takes the top 15 TFs by |Δedge_prob| (late−early) and e3 the top 15 TFs by maximum edge_prob, and when the same TF regulates multiple pathways the highest-scoring version is retained. **Sliding-window strategy.** For low-gene-number cases such as PVL and periodontitis, to increase statistical power each stage is enriched separately on 3 windows with 50% overlap, and significant terms are merged to reduce window-level stochastic fluctuation.

### 11. Evaluation metrics

**AUROC** measures edge-ranking quality. **Early precision ratio (EPR)** is the enrichment multiple of ground-truth edges among the top-N predictions (N = number of ground-truth positives).

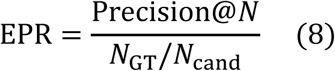

**Early count ratio (ECR)** is the multiple of the ground-truth edge count within top-N relative to the random expectation, measuring the gain in absolute top-hit count. **Directional accuracy (DirAcc)** is defined as

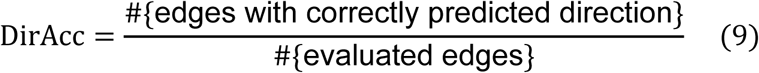

and is evaluated only on predicted edges with edge_prob > 0.2, reported separately for TF-to-non-TF (NT domain), TF-to-TF (TT domain), and all edges. **Confidence calibration.** Predicted edges are binned by edge_prob (bin center 0.10/0.50/0.90) and the empirical DirAcc per bin computed; perfect calibration corresponds to DirAcc = model confidence (the y=x line), and τ=0.2 splits edges into discarded (≤0.2) and retained (>0.2) classes.

### 12. Statistical analyses

#### Differential expression

DEGs for each disease stage against its baseline are identified by Wilcoxon rank-sum test with Benjamini–Hochberg FDR correction (FDR < 0.05, |log2FC| > 0.5). **Perturb-seq ground truth.** HCT116 uses a Welch t test (p < 0.01, |log2FC| > 0.25); K562 follows the authors’ released Truth-seq calls (|z| ≥ 2.0). **Direction baseline controls.** Randomly permuting ground-truth edge directions and recomputing DirAcc serves as the null hypothesis (HCT116 0.501, K562 0.500, no systematic bias). **Multi-seed robustness.** The edge model and NT direction expert are each trained with four independent seeds, with inter-seed consistency reported (e.g., synthetic-data edge-prediction AUROC 0.867–0.870) and ensembled means reducing seed sensitivity. **GO enrichment significance.** NOM p and FDR q come from gseapy prerank GSEA; term-count comparisons (e.g., the max|NES| distribution difference between DEG-Expanded and the conventional strategy, p < 1e-100) use the Wilcoxon rank-sum test. **Ablation comparisons.** Paired configuration comparisons (e.g., P+D vs D alone, joint vs separated training, dual experts vs DAGMA) use identical BEELINE inputs and evaluation protocol.

### 13. Computational tools and software

Python 3.12; PyTorch 2.x (CUDA-accelerated training, CPU-supported inference); scikit-learn (Ledoit–Wolf precision matrix, evaluation metrics); SciPy (distance correlation, sparse matrices, Wilcoxon tests); gseapy (prerank GSEA); scanpy (h5ad reading, cell-annotation handling); h5py (h5ad/h5 data reading); matplotlib / numpy / pandas (plotting and data processing). Base dependencies are in requirements.txt and full-pipeline dependencies in requirements-pipeline.txt (pyproject.toml requires Python ≥ 3.10; the tested environment is 3.12). All scripts support the BOYUE_ROOT, BOYUE_DATA, and BOYUE_CKPT environment variables to override default paths.

## Data availability

BEELINE benchmark data come from the original benchmark release[25]. For Perturb-seq data, the HCT116 files come from the X-Atlas/Orion genome-wide screen[37] and the K562 files from the genome-wide CRISPRi screen of Replogle et al.[38]. Disease-case data come from NCBI GEO: NAFLD (GSE202379)[39], HCC (GSE149614)[43], periodontitis (GSE164241)[42], PVL (GSE196296)[44], and AD (GSE157827)[45]. The PVL healthy control is taken from the BM of the periodontitis dataset. All datasets are directly available from their respective public sources.

## Code availability

Reproducible code is publicly available at https://github.com/holaoctopus/BoYueGRN The code repository does not include figure-generation scripts; Trained model checkpoints (edge model 4 seeds, NT direction expert 4 seeds, TT direction expert, and 7 ablation models) are distributed via a HuggingFace model repository at https://huggingface.co/Holaoctopus/boyuegrn.

## Acknowledgements

This work was supported by the Sichuan Provincial Natural Science Foundation(2026NSFSC0666)

---

**Fig. S1.**
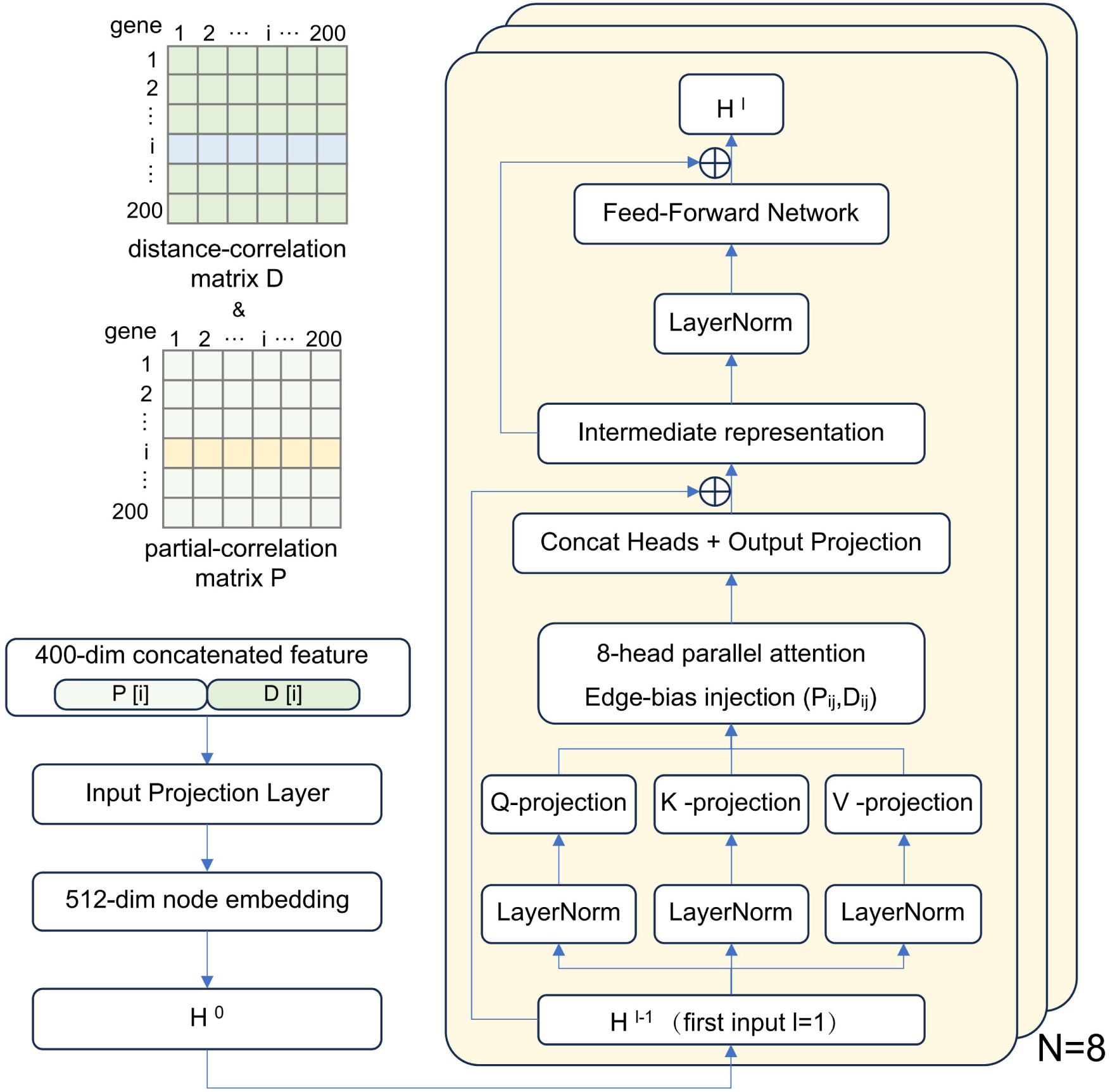
Input feature assembly and graph Transformer encoder architecture. Rows of the partial-correlation matrix P and the distance-correlation matrix D for each gene are concatenated into a 400-dimensional feature, projected by an input projection layer to a 512-dimensional node embedding (H⁰), and passed through N=8 identical encoder layers. Each layer applies LayerNorm, Q/K/V projections, 8-head parallel attention with edge-bias injection of the pairwise statistics (P_ij, D_ij) onto the attention logits, head concatenation with output projection, and a feed-forward network with residual connections.

**Fig. S2.**
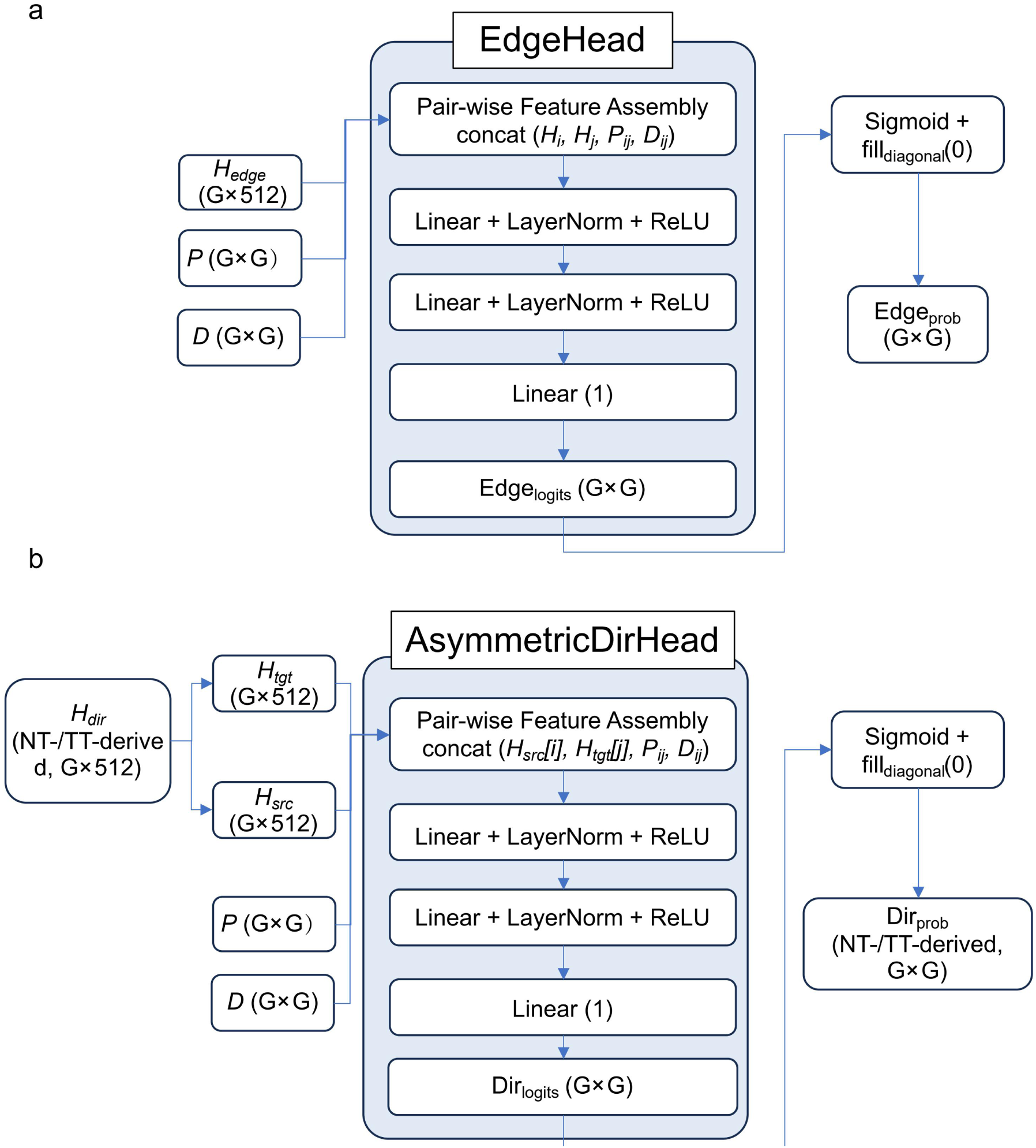
Prediction-head architectures. EdgeHead: pairwise feature assembly concatenates the two node embeddings with the corresponding P and D entries (concat(H_i, H_j, P_ij, D_ij)), followed by two Linear+LayerNorm+ReLU blocks, a scalar linear layer producing edge logits (G×G), and a sigmoid with zeroed diagonal yielding edge probabilities. AsymmetricDirHead: the direction encoder embedding is split into source and target projections (H_src, H_tgt); pairwise features concat(H_src[i], H_tgt[j], P_ij, D_ij) pass through the same MLP structure to produce direction logits and, after sigmoid, direction probabilities for the NT- and TT-derived heads.

**Fig. S3.**
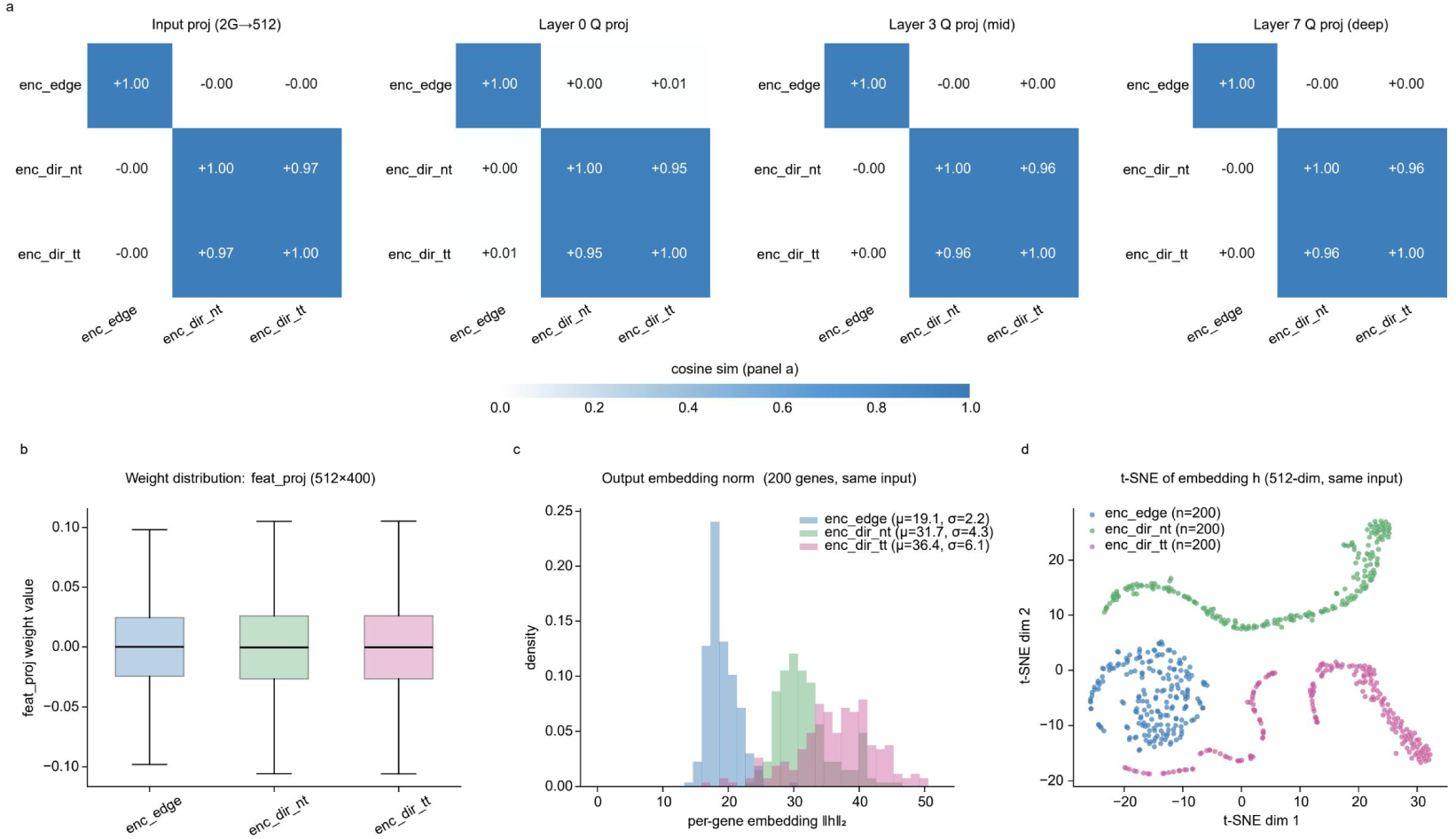
Differentiation of the independently trained encoders. a, Cosine similarity among the three encoders (enc_edge, enc_dir_nt, enc_dir_tt) at the input projection and at the Q projections of layers 0, 3, and 7: the two direction encoders remain similar to each other (about 0.95–0.98) while both diverge from the edge encoder (about 0). b, Weight distributions of the input feature projection (512×400) for the three encoders. c, Output embedding norms (200 genes, same input) for the three encoders (means 19.1, 31.7, and 36.4). d, t-SNE of the 512-dimensional output embeddings (same input), showing well-separated encoder-specific structure.

**Fig. S4.**
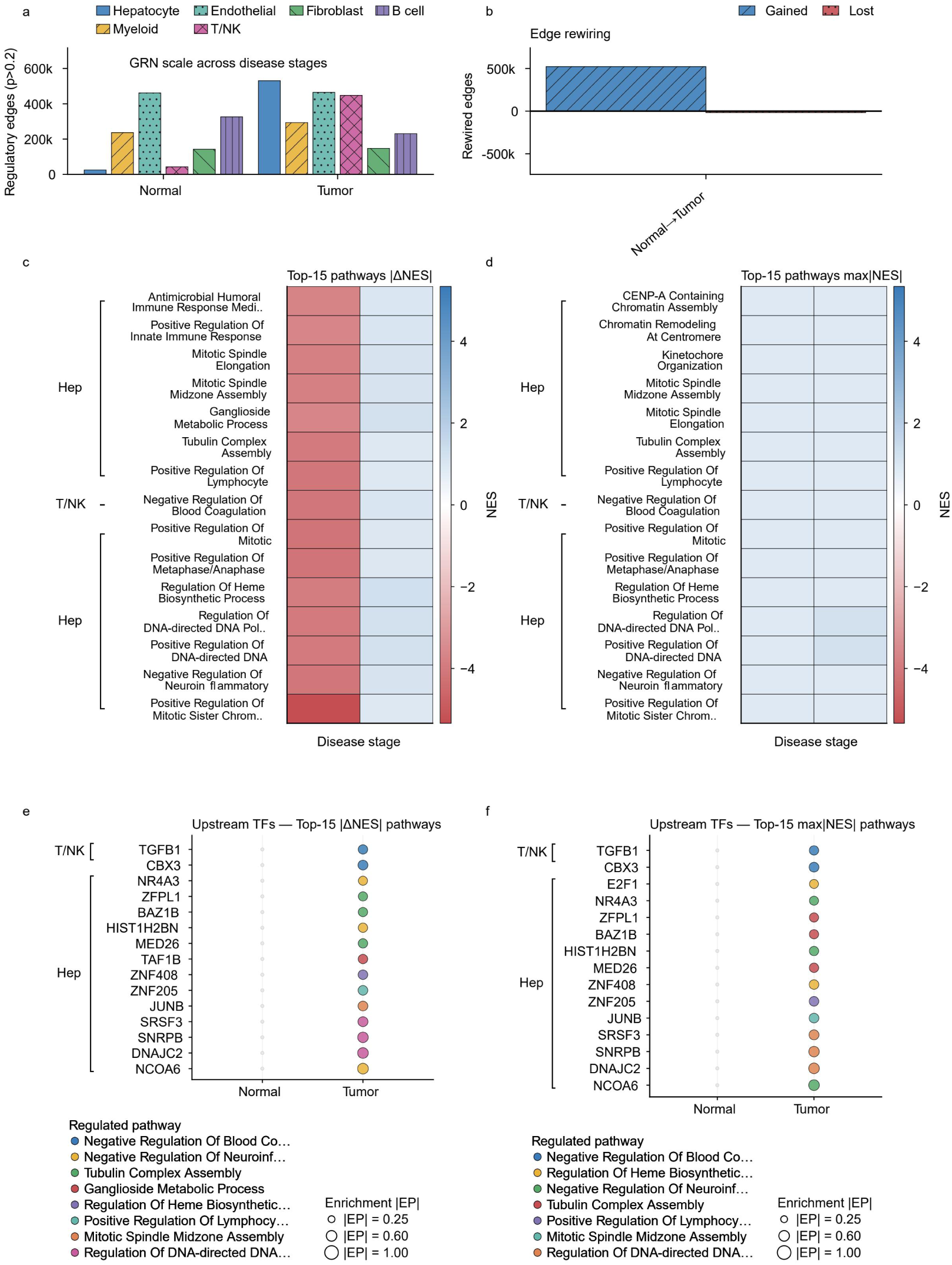
Hepatocellular carcinoma: network expansion and driver regulators. a, GRN scale in Normal versus Tumor liver across six cell types: the hepatocyte GRN expands 21-fold in an almost purely additive manner. b, Edge rewiring from Normal to Tumor (gained versus lost edges). c, d, Top-15 pathways ranked by |ΔNES| (c) and max|NES| (d), dominated by proliferative programs (mitotic spindle, CENP-A-containing chromatin assembly, kinetochore organization), consistent with a metabolic-to-proliferative transition. Per-cell-type TF-hub analysis (Supplementary Fig. S8) shows that MYC and E2F-family factors (E2F1, E2F3, E2F5) broaden their regulatory reach in the Tumor stage. e, f, Upstream regulators ranked by |ΔNES| (e) and max|NES| (f): NCOA6, DNAJC2, and SNRPB are returned independently by both rankings, pointing to metabolic reprogramming, ribosomal stress, and splicing dysregulation.

**Fig. S5.**
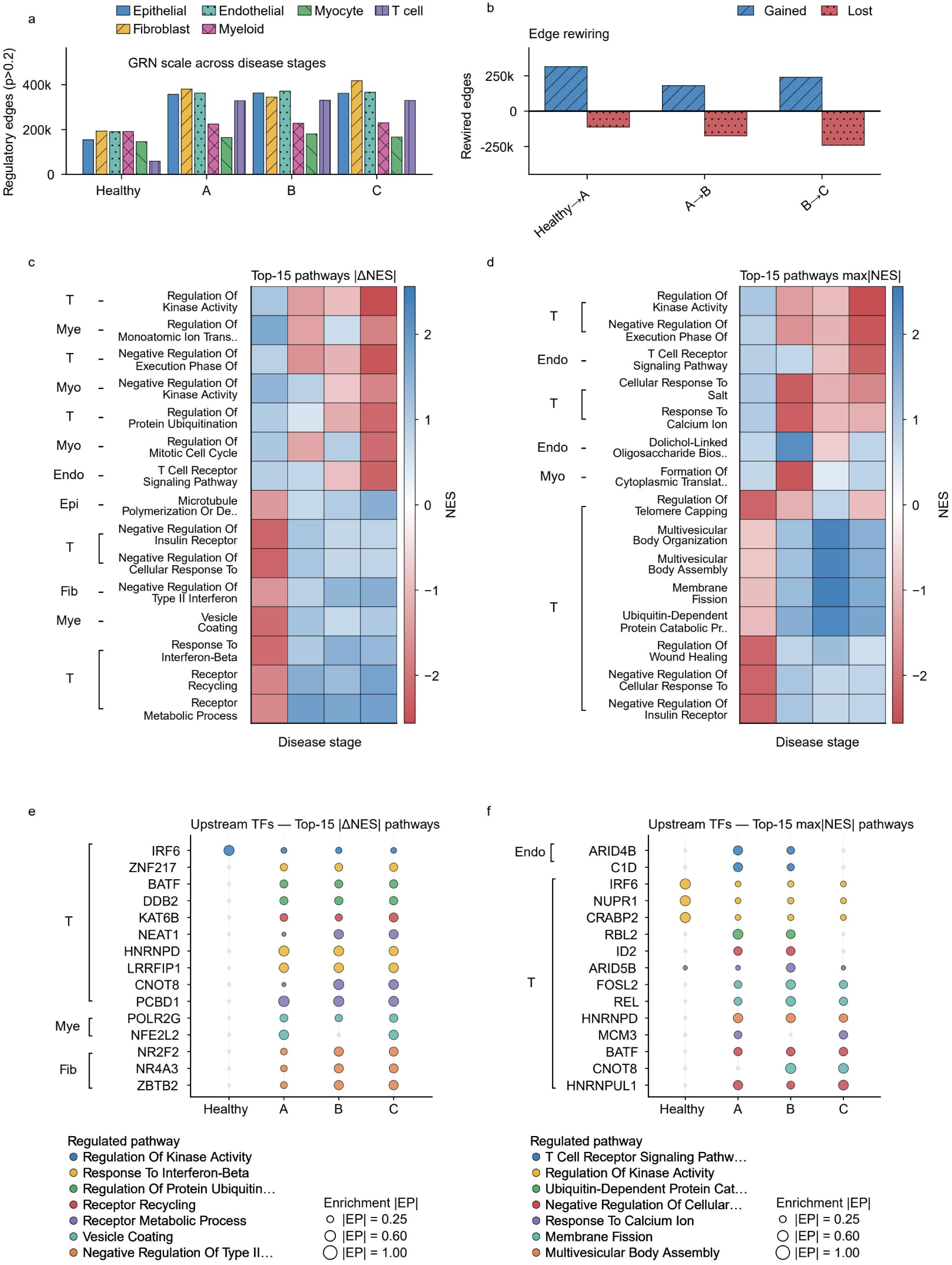
Proliferative verrucous leukoplakia: cell-type-heterogeneous regulatory responses. a, GRN scale across Healthy, A (mild), B (moderate), and C (severe) stages for six cell types: the T-cell GRN expands 5.6-fold and the epithelial GRN 2.3-fold. b, Edge rewiring across stage transitions. c, d, Top-15 pathways ranked by |ΔNES| (c) and max|NES| (d), with core perturbations in the microenvironmental compartment. e, f, Upstream regulators ranked by |ΔNES| (e) and max|NES| (f): driver reverse lookup independently recovers IRF6 (top-ranked in e), whose regulatory activity rises from mild to moderate stages (target genes expanding from 88 to 470).

**Fig. S6.**
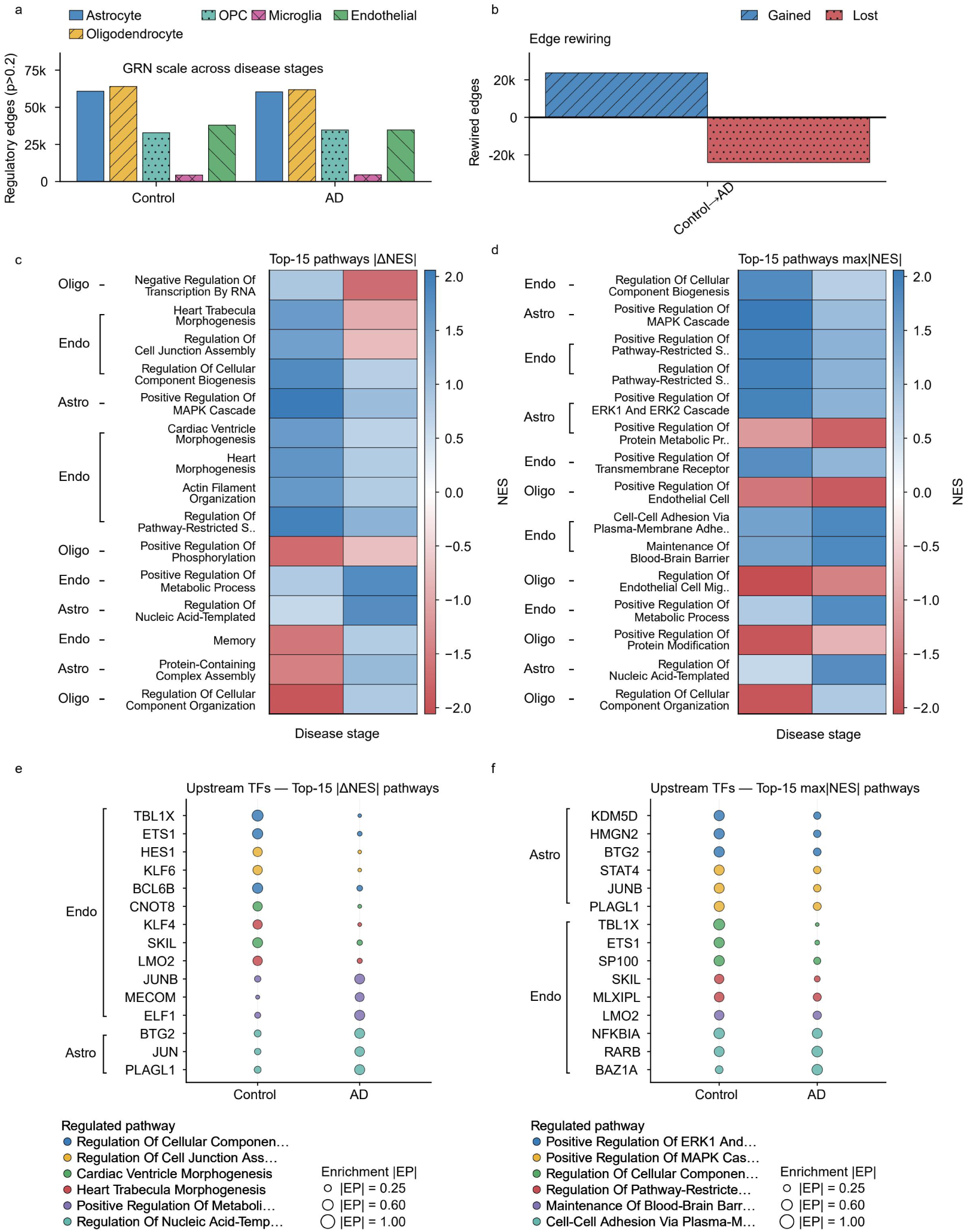
Alzheimer’s disease: mild network expansion with pathway-level changes in astrocytes. a, GRN scale in Control versus AD across five brain cell types (astrocytes, OPCs, microglia, endothelial cells, oligodendrocytes), showing mild expansion. b, Edge rewiring from Control to AD. c, d, Top-15 pathways ranked by |ΔNES| (c) and max|NES| (d): the most prominent astrocyte changes are enhancement of protein-containing complex assembly and protein-transport programs accompanied by weakening of MAPK/ERK cascade signaling, while oxidative phosphorylation, insulin-receptor signaling, and synaptic-organization terms appear but do not reach significance. e, f, Upstream regulators ranked by |ΔNES| (e) and max|NES| (f) across cell types.

## References

1. Mercatelli, D., et al., Gene regulatory network inference resources: A practical overview. Biochimica et Biophysica Acta (BBA)-Gene Regulatory Mechanisms, 2020. 1863(6): p. 194430.

2. Tang, F., et al., mRNA-Seq whole-transcriptome analysis of a single cell. Nature methods, 2009. 6(5): p. 377–382.

3. Badia-i-Mompel, P., et al., Gene regulatory network inference in the era of single-cell multi-omics. Nature Reviews Genetics, 2023. 24(11): p. 739–754.

4. Margolin, A.A., et al., ARACNE: an algorithm for the reconstruction of gene regulatory networks in a mammalian cellular context. BMC bioinformatics, 2006. 7(Suppl 1): p. S7.

5. Faith, J.J., et al., Large-scale mapping and validation of Escherichia coli transcriptional regulation from a compendium of expression profiles. PLoS biology, 2007. 5(1): p. e8.

6. Chan, T.E., M.P.H. Stumpf, and A.C. Babtie, Gene regulatory network inference from single-cell data using multivariate information measures. Cell systems, 2017. 5(3): p. 251–267.

7. Huynh-Thu, V.A., et al., Inferring regulatory networks from expression data using tree-based methods. PloS one, 2010. 5(9): p. e12776.

8. Moerman, T., et al., GRNBoost2 and Arboreto: efficient and scalable inference of gene regulatory networks. Bioinformatics, 2019. 35(12): p. 2159–2161.

9. Friedman, J., T. Hastie, and R. Tibshirani, Sparse inverse covariance estimation with the graphical lasso. Biostatistics, 2008. 9(3): p. 432–441.

10. Opgen-Rhein, R. and K. Strimmer, From correlation to causation networks: a simple approximate learning algorithm and its application to high-dimensional plant gene expression data. BMC systems biology, 2007. 1(1): p. 37.

11. Aibar, S., et al., SCENIC: single-cell regulatory network inference and clustering. Nature Methods, 2017. 14(11): p. 1083–1086.

12. Bravo González-Blas, C., et al., SCENIC+: single-cell multiomic inference of enhancers and gene regulatory networks. Nature Methods, 2023. 20(9): p. 1355–1367.

13. Kamimoto, K., et al., Dissecting cell identity via network inference and in silico gene perturbation. Nature, 2023. 614(7949): p. 742–751.

14. Kartha, V.K., et al., Functional inference of gene regulation using single-cell multi-omics. Cell Genomics, 2022. 2(9): p. 100166.

15. Yuan, Q. and Z. Duren, Inferring gene regulatory networks from single-cell multiome data using atlas-scale external data. Nature Biotechnology, 2024. 43(2): p. 247–257.

16. Yuan, Y. and Z. Bar-Joseph, Deep learning for inferring gene relationships from single-cell expression data. Proceedings of the National Academy of Sciences, 2019. 116(52): p. 27151–27158.

17. Deshpande, A., et al., Network inference with Granger causality ensembles on single-cell transcriptomics. Cell Reports, 2022. 38(6): p. 110333.

18. Shu, H., et al., Modeling gene regulatory networks using neural network architectures. Nature Computational Science, 2021. 1(7): p. 491–501.

19. Keyl, P., et al., Single-cell gene regulatory network prediction by explainable AI. Nucleic Acids Research, 2023. 51(4): p. e20–e20.

20. Theodoris, C.V., et al., Transfer learning enables predictions in network biology. Nature, 2023. 618(7966): p. 616–624.

21. Cui, H., et al., scGPT: toward building a foundation model for single-cell multi-omics using generative AI. Nature Methods, 2024. 21(**8****): p.** 1470**-**1480.

22. Hu, L., et al., RegFormer: a single-cell foundation model powered by gene regulatory hierarchies. Nature Communications, 2026. 17**: p.** 6432.

23. Marbach, D., et al., Revealing strengths and weaknesses of methods for gene network inference. Proceedings of the National Academy of Sciences, 2010. 107(14): p. 6286–6291.

24. Marbach, D., et al., Wisdom of crowds for robust gene network inference. Nature Methods, 2012. 9(8): p. 796–804.

25. Pratapa, A., et al., Benchmarking algorithms for gene regulatory network inference from single-cell transcriptomic data. Nature Methods, 2020. 17(2): p. 147–154.

26. Shimizu, S., et al., A linear non-Gaussian acyclic model for causal discovery. Journal of Machine Learning Research, 2006. 7: p. 2003–2030.

27. Hoyer, P.O., et al., Nonlinear causal discovery with additive noise models, in Advances in Neural Information Processing Systems 21 (NIPS 2008). 2008.

28. Zheng, X., et al., DAGs with NO TEARS: Continuous Optimization for Structure Learning. 2018, arXiv.

29. Löwe, S., et al., Amortized Causal Discovery: Learning to Infer Causal Graphs from Time-Series Data, in Conference on Causal Learning and Reasoning (CLeaR). 2022, arXiv.

30. Lorch, L., et al., Amortized Inference for Causal Structure Learning, in Advances in Neural Information Processing Systems 35. 2022, Neural Information Processing Systems Foundation, Inc. (NeurIPS). p. 13104–13118.

31. Dibaeinia, P. and S. Sinha, SERGIO: A Single-Cell Expression Simulator Guided by Gene Regulatory Networks. Cell Systems, 2020. 11(3): p. 252–271.e11.

32. Qiao, J., et al., CDFM: Towards a General-Purpose Causal Discovery Foundation Model. 2026, arXiv.

33. Tobin, J., et al., Domain randomization for transferring deep neural networks from simulation to the real world, in 2017 IEEE/RSJ International Conference on Intelligent Robots and Systems (IROS). 2017, IEEE. p. 23–30.

34. Barabási, A.-L.s. and R.k. Albert, Emergence of Scaling in Random Networks. Science, 1999. 286(5439): p. 509–512.

35. Ledoit, O. and M. Wolf, A well-conditioned estimator for large-dimensional covariance matrices. Journal of Multivariate Analysis, 2004. 88(2): p. 365–411.

36. Székely, G.J., M.L. Rizzo, and N.K. Bakirov, Measuring and testing dependence by correlation of distances. The Annals of Statistics, 2007. 35(6).

37. Huang, A.C., et al., X-Atlas/Orion: Genome-wide Perturb-seq Datasets via a Scalable Fix-Cryopreserve Platform for Training Dose-Dependent Biological Foundation Models. 2025, openRxiv.

38. Replogle, J.M., et al., Mapping information-rich genotype-phenotype landscapes with genome-scale Perturb-seq. Cell, 2022. 185(14): p. 2559–2575.e28.

39. Gribben, C., et al., Acquisition of epithelial plasticity in human chronic liver disease. Nature, 2024. 630(8015): p. 166–173.

40. Subramanian, A., et al., Gene set enrichment analysis: A knowledge-based approach for interpreting genome-wide expression profiles. Proceedings of the National Academy of Sciences, 2005. 102(43): p. 15545–15550.

41. Fang, Z., X. Liu, and G. Peltz, GSEApy: a comprehensive package for performing gene set enrichment analysis in Python. Bioinformatics, 2023. 39(1).

42. Williams, D.W., et al., Human oral mucosa cell atlas reveals a stromal-neutrophil axis regulating tissue immunity. Cell, 2021. 184(15): p. 4090–4104.e15.

43. Lu, Y., et al., A single-cell atlas of the multicellular ecosystem of primary and metastatic hepatocellular carcinoma. Nature Communications, 2022. 13(1).

44. Zhong, L., et al., Single-cell transcriptomics dissects premalignant progression in proliferative verrucous leukoplakia. Oral Diseases, 2024. 30(2): p. 172–186.

45. Lau, S.-F., et al., Single-nucleus transcriptome analysis reveals dysregulation of angiogenic endothelial cells and neuroprotective glia in Alzheimer’s disease. Proceedings of the National Academy of Sciences, 2020. 117(41): p. 25800–25809.

46. Moya, I.M. and G. Halder, Hippo–YAP/TAZ signalling in organ regeneration and regenerative medicine. Nature Reviews Molecular Cell Biology, 2019. 20(4): p. 211–226.

47. Botti, E., et al., Developmental factor IRF6 exhibits tumor suppressor activity in squamous cell carcinomas. Proceedings of the National Academy of Sciences, 2011. 108(33): p. 13710–13715.

48. Bello, K., B. Aragam, and P. Ravikumar, DAGMA: Learning DAGs via M-matrices and a Log-Determinant Acyclicity Characterization. 2022, arXiv.

49. Vaswani, A., et al., Attention Is All You Need. 2017, arXiv.

50. Kingma, D.P. and J. Ba, Adam: A Method for Stochastic Optimization. 2014, arXiv.

